# Mice learn multi-step routes by memorizing subgoal locations

**DOI:** 10.1101/2020.08.19.256867

**Authors:** Philip Shamash, Sarah F. Olesen, Panagiota Iordanidou, Dario Campagner, Banerjee Nabhojit, Tiago Branco

## Abstract

The behavioral strategies that mammals use to learn multi-step routes in natural settings are unknown. Here we show that mice spontaneously adopt a subgoal memory strategy. We first investigated how mice navigate to shelter in response to threats when the direct path is blocked. Initially, they fled toward the shelter and negotiated obstacles using sensory cues. Within twenty minutes, they adopted a subgoal strategy, initiating escapes by running directly to the obstacle’s edge. Mice continued to target this subgoal location after the obstacle was removed, indicating use of spatial memory. However, standard models of spatial learning – egocentric-movement repetition and internal-map building – did not explain how subgoal memories formed. Instead, mice used a hybrid approach: memorizing salient locations encountered during spontaneous ‘practice runs’. This strategy was also used during geometrically identical reward-seeking behavior. These results suggest that subgoal memorization is a fundamental strategy by which rodents learn efficient multi-step routes in new environments.

## Introduction

For prey species such as mice, quickly finding effective routes to goals is critical for survival because it reduces exposure to potential predators (Lima and Dill, 1990). This is a challenging task: natural environments are complex, and wild animals must compute multi-step routes taking into account uneven terrain, obstacles, and dynamically changing environments. Ethological studies of wild rodents have emphasized the roles of locating salient landmarks (Drickamer and Stuart, 1984; McMillan and Kaufman, 1995) and adhering to familiar paths (Benhamou, 1991; Thompson, 1982) in overcoming these challenges. However, observational studies are limited in their ability to identify the cues and behavioral strategies that animals actually use to navigate.

Experimental evidence from rodents trained to locate goals has uncovered multiple types of spatial reasoning that can be used to solve complex navigational problems. On the one hand, rodents keep track of their position within an allocentric (environment-centered) reference frame (Morris, 1981). This sense of place is thought to be integrated into an internal topological map connecting locations within the environment, which allows animals to compute subgoal locations whenever a new multi-step route to a goal is required (Edvardsen et al., 2020; Spiers and Gilbert, 2015; Stachenfeld et al., 2017; Tolman, 1948). This kind of cognitive-map-based reasoning is flexible, is learned by observing the structure of the environment, and it depends on the hippocampus (O’Keefe and Nadel, 1978). Alternatively, animals can navigate to goals without relying on an internal map. These strategies include integrating self-motion cues to compute a vector back to their starting position (Etienne and Jeffery, 2004); repeating egocentric movements at familiar junctions (Restle, 1957; Sutherland and Dyck, 1984); and using landmarks for visual guidance (Hamilton et al., 2004). The latter two tactics, known as “taxon” strategies, are inflexible, rely on proximal cues, and are learned through previous motivated actions (O’Keefe and Nadel, 1978).

Despite all that is known about rodent navigation, the behavioral strategies that animals spontaneously use to quickly build up and deploy spatial knowledge in new environments remain unknown. The abilities listed above have mostly been demonstrated by repeatedly placing rodents in constrained mazes until they learn to navigate to a goal. In a natural setting, however, spatial learning must occur via internally generated exploration patterns and within a very limited timeframe. It is therefore unclear how well previous classifications of navigation strategies map onto the instincts and learning procedures that animals use during natural goal-directed navigation.

Escape behavior offers a powerful model for studying naturalistic navigation in the laboratory. Diverse animals, including fishes, lizards, crabs, birds, and rodents, respond to threats by escaping to a familiar shelter (Cooper Jr. and Blumstein, 2015). Mice are known to rapidly identify and memorize shelter locations in new environments and instinctively respond to visual or auditory threats by running straight to the shelter (Vale et al., 2017; Yilmaz and Meister, 2013). Previous studies have shown that the spatial memory for running back to shelter (‘homing’) can be based on path integration or distal visual landmarks when a direct path is available. (Alyan and Jander, 1994; Etienne et al., 1985; Harrison et al., 2006; Vale et al., 2017). If the direct path is blocked on one side by a barrier, previous work has shown that gerbils can use spatial memory to reach the hidden shelter after a brief period of exploration (Ellard and Eller, 2009). Thus, rodent escape offers not only reliable, stimulus-locked trajectories and rapid learning within a single session but also a reliance on spatial reasoning. These qualities make escape a useful model for understanding how animals learn and execute complex goal-directed trajectories within the time constraints compatible with survival in natural settings.

Here we first investigate the strategies that naïve mice use to navigate to shelter in response to auditory threats when the direct path is blocked by a wall. Through quantitative analysis of escape trajectories and their relationship to exploratory behavior, systematic variation of spatial conditions, and dynamic modifications to the environment, we describe how mice learn to execute efficient multi-step escape routes within minutes of entering a novel, obstacle-laden environment. We then show that the navigational strategy for escape is also used for reaching a food reward goal.

## Results

### Mice rapidly learn efficient escape routes in the presence of an obstacle

As a baseline condition for investigating how mice learn escape trajectories, we placed naïve animals in a circular, open-field platform with a shelter and overhead lighting. After a brief exploration period during which mice spontaneously located the shelter, we exposed them to a loud, overhead crashing sound while they were in a pre-defined threat zone (Fig. 1A). This reliably elicited rapid escapes directed at the shelter along a straight ‘homing vector’ (N=23 escapes, 10 mice; Fig. 1A-B, Extended Data Fig. 2A; Supplementary Video 1-2), similar to previous results (Vale et al., 2017). We then repeated this experiment in a separate group of mice, with a wall positioned between the threat zone and the shelter (N = 24 mice; Extended Data Fig. 1A). This wall was white against a black background, and all mice approached and walked along it during the exploration period (Extended Data Fig. 2B). To quantify escape trajectories in relation to the obstacle, we computed a target score: escapes aimed at the shelter get a score of zero; escapes targeting the obstacle edge get 1.0; and escapes aimed beyond the obstacle edge get scores >1.0 (Fig. 1A). Escapes are classified as “edge vectors” if their score surpasses the 95^th^ percentile of escape scores in the open field (0.65) and are otherwise classified as “homing vectors”. Upon the first threat presentation, the majority of the mice (57%) executed homing-vector escapes (Fig. 1A-B; Supplementary Video 3). Replacing the wall obstacle with an unprotective hole obstacle did not reduce this proportion (Extended Data Fig. 1B, Extended Data Fig. 2C-D); thus, homing-vector escapes cannot be accounted for by the safety provided by running along a wall and are likely directed at the shelter location.

**Figure 1.**
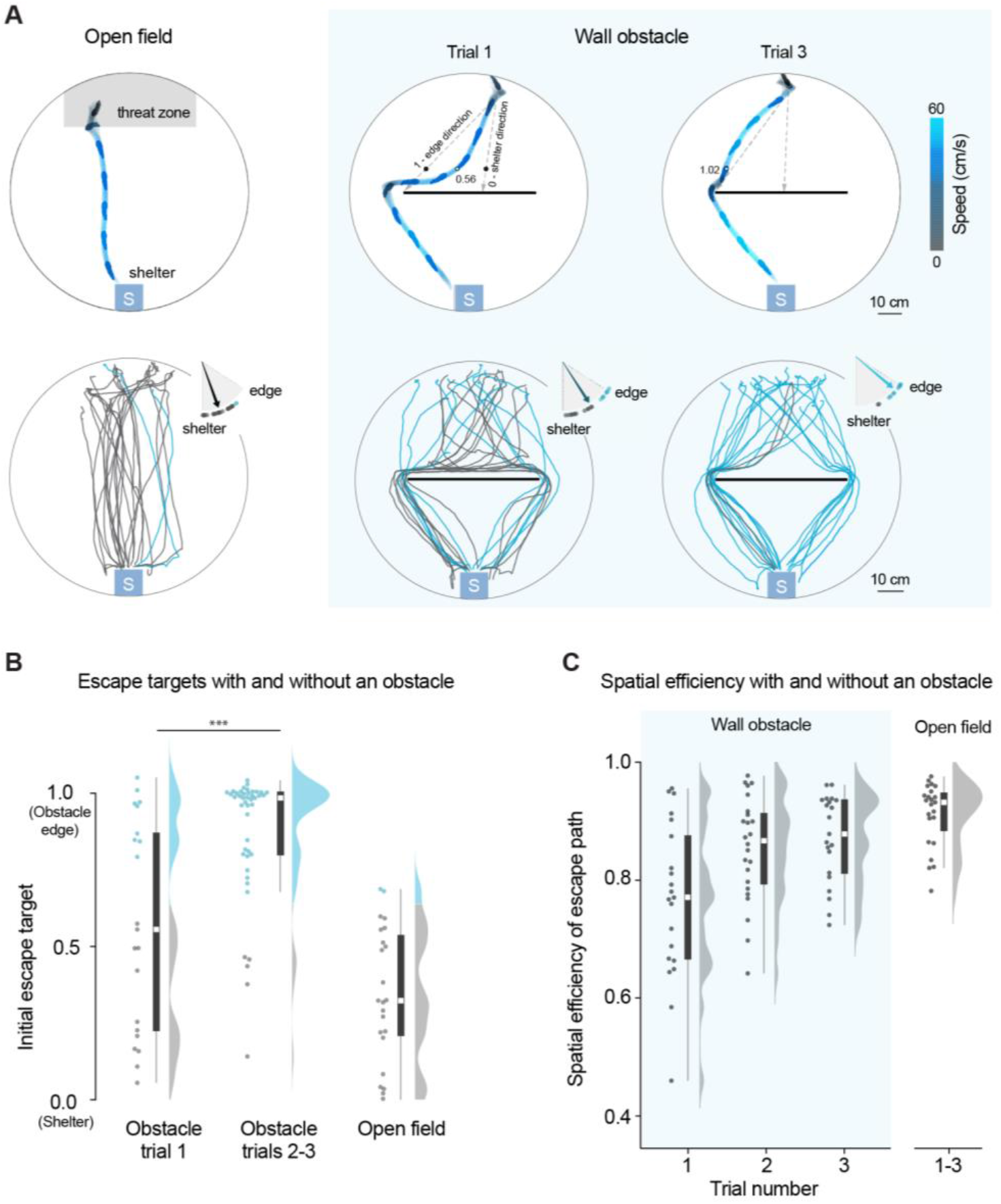
Mice rapidly learn efficient escape trajectories in the presence of obstacles. (A) Single escape trials colored by speed (top) and all trajectories color-coded by trajectory type (bottom; homing-vector paths are gray, edge-vector paths are blue). Initial escape target is the direction of the vector from escape initiation to 10 cm in front of the obstacle, normalized between 0 (directed at the shelter) and 1 (directed at the obstacle edge location). Dashed arrows in the top panels illustrate the shelter and edge-vector directions. Dots show the point where the target measurement is taken. Values (0.56 and 1.02) are the escape target score for the examples shown. Dot-and-arrow plots show the distribution of initial escape targets (the arrow is the median value). Only successful escapes (getting to the shelter within 12 seconds) are analyzed. Obstacle: trial 1: 21 escapes; trial 2: 24 escapes; trial 3: 21 escapes. Total: N=66 escapes, 24 mice. Open field total: N=23 escapes, 10 mice. (B) Summary of initial escape targets. Permutation test on the target score, trial 1 vs. trials 2-3: P=0.0004 (***). Each dot represents one escape (2-3 escapes per mouse). Gray points are escapes classified as homing vectors and blue points are edge-vector escapes. White squares show the median, thick lines show the IQR, and thin lines show the range excluding outliers. Distributions are kernel density estimates. Differences in escape target were not related to differences in position or angle at the moment of threat onset (see Extended Data Fig. 10). (C) Spatial efficiency is the ratio of the shortest possible escape path length and the path actually taken to the shelter. This is the inverse of tortuosity. Each dot represents one escape.

Over the course of three threat presentation trials (17±4 minutes into the session, *mean±std*), mice performed escapes that were increasingly spatially efficient (ratio of the shortest possible path to the actual escape path: median for trial 1 = 0.77; for trial 3 = 0.87; F(2, 30)=7.2, P=.003, *repeated measures ANOVA on trials 1-3;* Fig. 1C) and rapid (normalized escape duration: median for trial 1 = 3.8 s; for trial 3 = 3.2 s; F(2, 34)=6.2, P=.005; Extended Data Fig. 2E). By this point, almost all trajectories were aimed directly at the obstacle edge (90% edge vectors; median target score = 0.98). Thus, while inefficient homing responses initially dominated, mice acquired rapid and streamlined routes to shelter over the course of 20 minutes and three escape trials.

We next investigated whether mice use visual input to locate and run toward the obstacle edge. First, we examined how mice responded to an unexpected obstacle rising up at the same time as the threat stimulus onset (N=10 trials, 10 mice); this trial occurred following 20 minutes with three escape trials in the open field. We found that 7/10 escapes followed homing-vector paths until reaching the wall. The other 3/10 escapes deviated toward an obstacle edge before they were close enough to touch the wall, indicating that vision was used to navigate toward the obstacle edge (putative visual obstacle avoidance; 3/10 > 0/23 escapes in the open field; P=0.02, *Fisher’s exact test;* Extended Data Fig. 3A; Supplementary Video 4). This suggests that visual and tactile cues can both be used to negotiate obstacles during escape, in the absence any experience with an obstacle. Second, we examined whether visual cues were *necessary* for generating edge-vector escapes. We repeated the obstacle experiment from Figure 1, but now in complete darkness (Extended Data Fig. 3B-D; Supplementary Video 5). Mice now executed fewer edge-vector escapes (% edge-vector escapes on trials 1-3: 33% with the lights off vs. 74% with the lights on, P=0.002, *permutation test*), to a level that was not significantly different than chance (P=0.2 for the comparison with the 22% edge-vector escapes in the dark without an obstacle; *permutation test*). However, after 20 minutes with three escape trials in the light, mice were able to execute mostly edge vectors in the dark (55% edge-vector escapes vs. 22% in the open field, P=0.002, *permutation test;* Extended Data Fig. 3C-D). Thus, for naïve mice with limited experience, visual cues are required for efficient obstacle avoidance. However, immediately after experiencing a 20-minute behavioral session, streamlined escapes can occur even in complete darkness.

### Mice develop a spatial memory strategy for efficient obstacle avoidance

We thus considered that learning efficient escapes might entail developing a memory of the obstacle edge location, making perception of the obstacle unnecessary. To further test this hypothesis, after the animals explored the environment with the obstacle for 20 minutes and with three escape trials, we removed the obstacle at the moment of threat onset (“acute obstacle removal”; Supplementary Video 6). Although the obstacle disappeared before the initial orientation movement could be completed, all animals escaped along the edge vector and did not turn toward the shelter until they passed the location where the obstacle edge used to be (median target = 0.98; N=8 escapes, 8 mice; Fig. 2A-B). Next, we examined how persistent this memory-based strategy is. In a “chronic obstacle removal” experiment (CORE), we allowed mice to explore after this acute obstacle removal trial (for 9±5 minutes, *mean±std*), during which time they visited the now empty center of the platform (Extended Data Fig. 4A). 44% of the subsequent escapes were still directed at the location where the obstacle edge used to be (N=18 escapes, 8 mice; more than the 9% edge-vector rate in the open field: P=0.02, *permutation test;* Fig. 2A-B; Supplementary Video 6), while the remaining 56% mice reverted to the homing-vector response.

**Figure 2.**
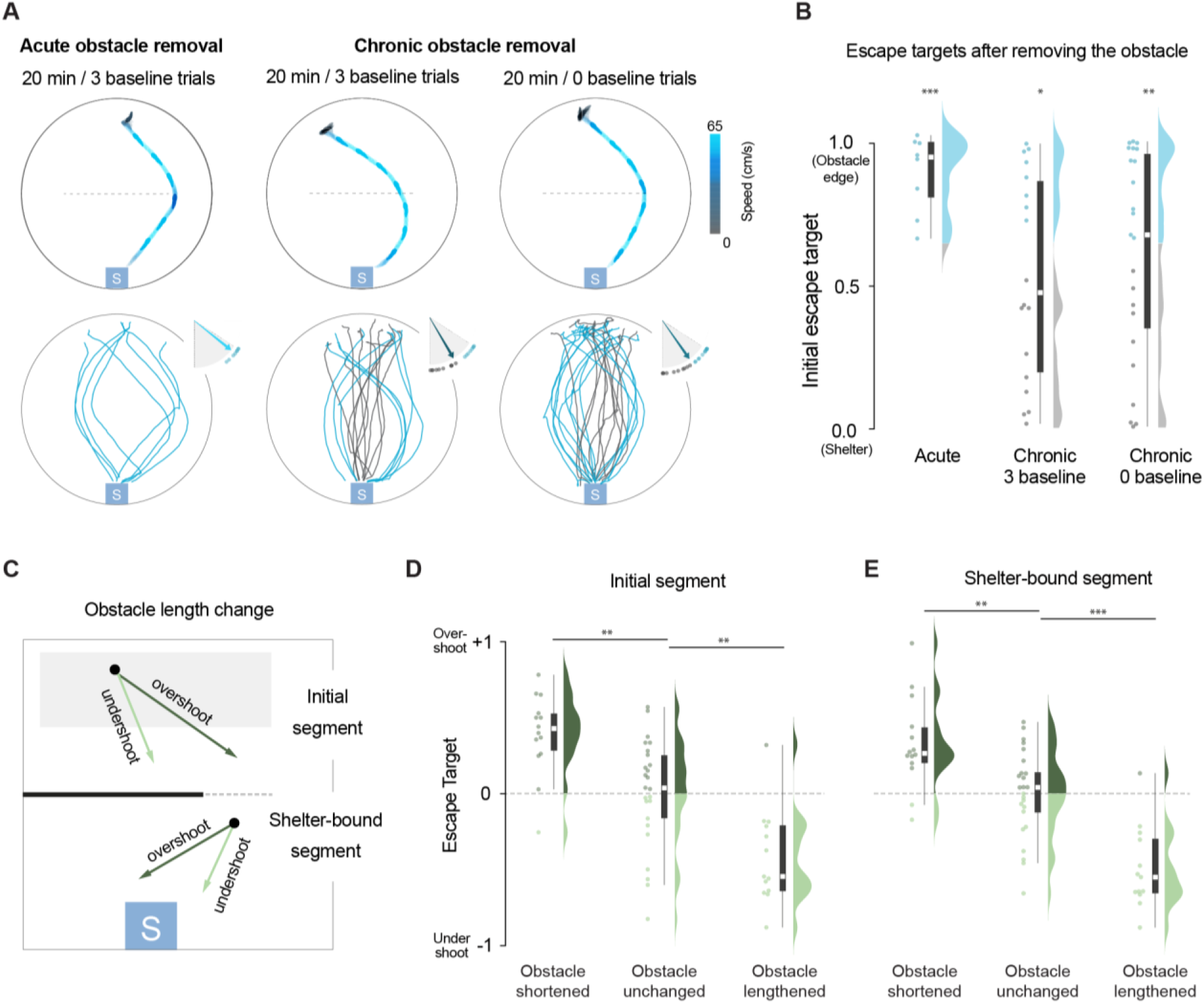
Mice use a spatial memory strategy for efficient obstacle avoidance. **(A)** Examples of edge-vector escape trials (top) and all trajectories (bottom) after removing the obstacle. Subheaders describe the experience in the environment prior to removing the obstacle. The dotted line indicates where the obstacle used to be. Arrow plots show the distribution of initial escape targets. Acute removal: N=8 escapes, 8 mice; chronic removal, 3 baselines: N= 18 escapes, 8 mice; chronic removal, 0 baselines: N=23 escapes, 10 mice. **(B)** Summary data for initial escape targets after obstacle removal. * P<0.05, ** P<0.01, *** P<0.001, permutation test on the rate of edge-vectors compared to the open field. (C) Schematic for the experiment in which the obstacle length is changed by 25%. For the “obstacle shortened’’ condition, the dotted gray line indicates the initial obstacle length, and the thick black line indicates the new length (after 20 minutes / 3 baseline escape trials). For the “obstacle lengthened” condition, the thick black line indicates the initial obstacle length, and the dotted gray line indicates the new length. Both the initial and the shelter-bound segments of the escape are analyzed. (D) Summary data for the experiments changing the length of the obstacle by 25%, analyzing the initial segment of the escape. Escapes aimed at the current obstacle edge location get a score of zero; escapes aimed at the edge location if the obstacle were 25% longer get a score of +1 (overshoot); and escapes escapes aimed at the edge location if the obstacle were 25% shorter get a score of −1 (undershoot). Combining both length-change experiments, the median of the initial target score’s absolute value is 0.44. Permutation test on the initial escape target, obstacle shortened vs. unchanged: P=0.002; obstacle lengthened vs. unchanged: P=0.002. (E) Summary data for the experiments changing the length of the obstacle by 25%, analyzing the segment of the escape going from the obstacle edge to the shelter. Escapes aimed at the current shelter location get a score of zero; escapes aimed where the shelter would be if the obstacle were 25% longer get a score of +1 (overshoot); and escapes aimed where the shelter would be if the obstacle were 25% shorter get a score of −1 (undershoot). Combining both length-change experiments, the absolute median shelter-bound target score magnitude is 0.37. Permutation test on the shelter-bound escape target, obstacle shortened vs. unchanged: P=0.001; obstacle lengthened vs. unchanged: P=0.0002.

This spatial memory for edge-vector escapes could in principle be learned during escapes trials or through spontaneous exploratory behavior. To distinguish between these possibilities, we repeated the CORE with zero threat stimuli during the initial 20-minute exploration period. As in the previous experiment, we then removed the obstacle and allowed the mice to explore the newly unobstructed environment (for 5±4 minutes, *mean±std*). Threat presentation after this period resulted in mostly edge-vector responses (57% edge-vector escapes; N = 23 escapes, 10 mice; more edge vectors than in the open field: P=0.004, *permutation test;* Fig. 2A-B). Thus, within 20 minutes in a novel environment, mice spontaneously develop a persistent spatial memory for efficient, multi-step escapes.

Our experiments so far have revealed a memory-based obstacle avoidance strategy. We have also shown, however, that visual input is used to navigate around the obstacle when experience is limited. We therefore tested how perception and spatial memory operate in tandem when both are fully available. In a novel environment, we performed an experiment similar to the CORE, but instead of removing the obstacle, we changed its length by 25% (Fig.2C; obstacle shortened: N=14 escapes, 9 mice; obstacle lengthened: N=13 escapes, 9 mice; obstacle always short: N=15 escapes, 9 mice; obstacle always long: N=10 escapes, 8 mice). Initial escape trajectories were consistently biased toward the previous edge location (Fig. 2D-E, Extended Data Fig. 4B-E; Supplementary Video 7). However, this result differed from the obstacle removal experiments in two ways. First, the memory bias was intermediate in magnitude: escape targets were biased only partway toward the former edge location (Fig. 2D, Extended Data Fig. 4D-E). Second, the second segment of the escape was equally biased: after reaching the obstacle edge, mice ran toward where the shelter would be if the edge had not moved (Fig. 2E, Extended Data Fig. 4D-E). These results show that, when available, the current obstacle position is an important cue for anchoring and adjusting memory-guided paths to both the edge and the shelter.

### Characterizing the spatial memory strategy for escape past an obstacle

We next aimed to characterize the spatial-memory strategy and how it is learned. We evaluated three possible strategies: habitual learning of turn angles, sampling the environment to build a cognitive map, and memorizing subgoals encountered during practice homings. We evaluated each possibility by analyzing the relationship between escapes and spontaneous behavior during exploration, primarily in the chronic obstacle removal experiment with zero baseline trials (CORE-ZB). For each analytical finding, we then performed further experiments to validate the analysis.

#### Habitual, egocentric movements do not explain the spatial memory for escape: analysis

First, we tested whether mice learn egocentric movements from the threat zone to the obstacle edge, similar to the habitual response strategy in mazes (Restle, 1957). We extracted all spontaneous homing runs, defined as sustained turn-and-run movements from the threat area toward the shelter during the CORE-ZB’s exploration period (see Methods; median [IQR] number of runs = 7 [6,7]; time from their end point until reaching the shelter: 11 [5, 19] sec; Fig. 3A; Extended Data Fig. 5A; Supplementary Video 8). We then computed each run’s starting position and orientation, and the angle turned during its initial turn-and-run segment (Fig. 3A; see Methods and Extended Data Fig. 10 for examples of starting-point extraction).

**Figure 3.**
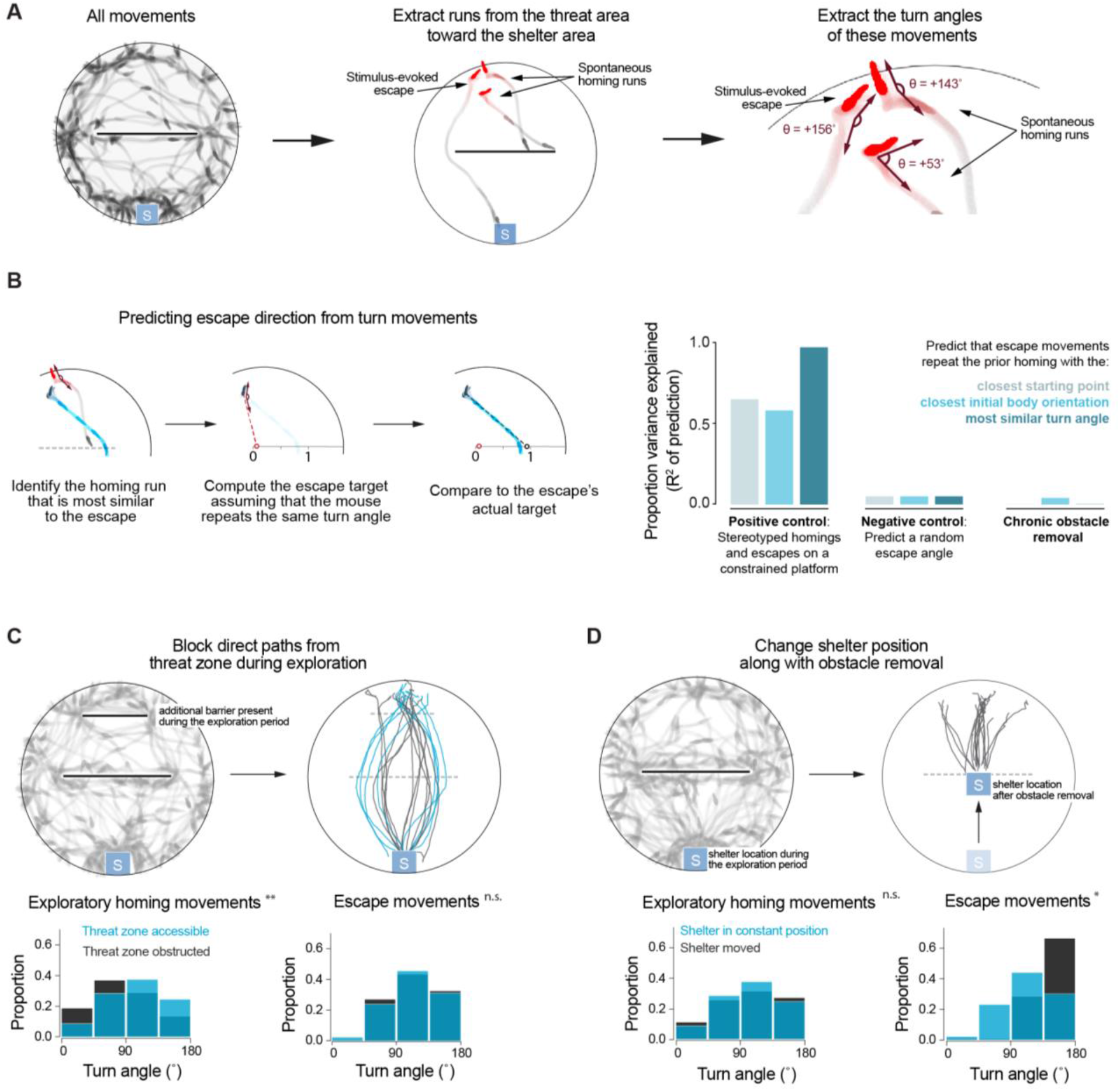
Habitual, egocentric movements do not explain the spatial memory for escape. **(A)** Examples of homing runs, continuous turn-and-run movements toward the shelter, extracted from all movement data (Supplementary Video 9; Methods). Turn angles are defined as the difference in heading direction between the homing initiation point and the point where the mouse has travelled 15 cm. Turn angles are positive for rightward turns and negative for leftward turns. (B) Predicting the escape targets of mice in the CORE-ZB on the basis of their previous turning movements. Left: diagram illustrating the method for predicting the escape target based on the turn angles from previous homing runs. Right: the y-a×is measures the amount of variance in escape targets across trials that can be explained by mice repeating turn angles from previous homing runs. The positive ∞ntrol is a platform with narrow corridors, in which movements are stereotyped (see Extended Data Fig. 5; N=30 escapes, 10 mice). The test condition is the CORE-ZB. The negative control is predicting a random angle, and then extrapolating that angle to predict escape targets in the CORE-ZB (avg R^2^ from doing this 1000 times). (C) An experiment similar to the CORE-ZB, but with an additional barrier present during exploration. N=20 escapes, 10 mice. Bottom: distribution of turn angles from homing movements and from escapes. Both sets of angles are extracted in the same way (Fig. 3a; Methods). Black shows the new CORE with two barriers, and blue shows the two original COREs (Fig. 2A). * P<0.05, ** P<0.01; chi-square test comparing the angle distributions, binned as shown in the figure. (D) An experiment like the CORE-ZB, but in which the shelter is moved following the exploration period. N=18 escapes, 10 mice. Analysis is the same as in panel C. Bottom: Black shows the new CORE with the moved shelter, and blue shows the two original CORES.

Homing runs were sparse, and their initial positions and body orientations were highly variable. It was unlikely for any escape’s starting conditions to closely match a previous homing: only 22% of escapes were preceded by a run with starting points within 10 cm distance and 30° body orientation (Extended Data Fig. 5A). Despite this lack of stereotypy, we attempted to account for the memory-guided edge-vector escapes observed in the CORE-ZB using the assumption that mice repeat turn angles from previous homing runs. First, we validated a method to predict escape targets based on homing-run turn angles. We put mice on a modified platform with two narrow corridors, ensuring that homings and escapes were stereotyped (N=30 escapes, 10 mice; Extended Data Fig. 5B). Here, we could precisely predict escape targets using the mouse’s starting point and its history of previous turn-and-run movements (R^2^ of the prediction = 0.97 using the homing run with the most similar turn angle; R^2^ = 0.65 using the homing run with the closest initial position; R^2^ = 0.58 using the homing run with the closest initial body orientation; Fig. 3B; Extended Data Fig. 5C). In the CORE-ZB, however, repeating turning movements did not explain any of the variance in post-removal escape targets (R^2^ of the prediction = 9×10^-4^ (most similar turn angle); R^2^ = 0.04 (closest initial position); R^2^ = 9×10^-6^ (closest initial body orientation); R^2^ = 0.05 (randomly generated prediction); Fig. 3B; Extended Data Fig. 5D).

#### Habitual, egocentric movements do not explain the spatial memory for escape: experiments

This analysis suggests that memory-guided edge-vector escapes are not based on repeating egocentric actions. We performed two new experiments to test this finding. First, we performed a variant of the CORE-ZB designed to induce a distinct pattern of movements during exploratory paths from the threat zone toward the shelter. We added an additional barrier in front of the threat zone, which was removed simultaneously with the main obstacle (Fig. 3C; N=20 escapes, 10 mice). This new geometry altered the homing-run turn angles during exploration (relative to the two COREs with the original geometry; P=0.001, *chi-square test on the distribution of homing-run turn angles;* Fig. 3C). However, the distribution of escape turn angles was indistinguishable (P=0.6, *chi-square test on the distribution of escape turn angles;* Fig. 3C). Thus, different fine-grained movements during exploration do not necessarily produce different escape movements. We also observed that escape routes did not deviate around the second, distal obstacle location (Fig. 3C). This further suggests that memory-guided escapes are not merely a function of the movements made during exploration but instead depend on the geometry of the mouse’s location, the obstacle, and the shelter. In a second experiment, we directly tested whether memory-guided escapes are sensitive to the shelter location; this would not be expected from a habitually repeated action (Fig. 3D; N=18 escapes, 10 mice). After a 20-minute exploration period just like in the CORE-ZB, we moved the shelter to the middle of the platform. As expected from a goal-directed or geometry-dependent process, escape turns differed from the two original COREs, with zero escapes targeting the obstacle edge location (P=0.5, *chi-square test on homing-run turn angles;* P=0.03, *chi-square test on escape turn angles;* Fig. 3D). Both of these experiments demonstrate a dissociation between egocentric turning movements and memory-guided escapes: distinct exploratory movements can lead to identical escape movements, and identical exploratory movements can lead to distinct escape movements.

#### Mice memorize previously encountered target locations as subgoals: analysis

The goal-directed nature of these escapes suggests that the obstacle edges become subgoal locations. An alternative possibility, however, is that mice target the edge by learning allocentric heading directions. For example, edge-vector escapes could be generated by consistently running in the southwest or southeast direction, relative to the north-south axis connecting the shelter and the threat zone. Analysis of our data indicates that mice instead target allocentric *locations*. Following obstacle removal, escape heading directions follow whichever direction is required to reach the edge location (correlation between the heading direction to the edge and the heading direction taken in the escape: r=0.85, P=1.2×10^-6^; Extended Data Fig. 5E). This corroborates the results above, suggesting that mice learn true subgoals at the obstacle edge.

We next investigated the learning process that generates these subgoals during the spontaneous exploration period. We found two variables in the CORE-ZB with high, positive correlations to subgoal-targeting behavior: the total distance of exploratory movement on the threat side of the platform (correlation with post-removal escape targets: r=0.72, P=1×10^-4^; Extended Data Fig. 6A) and the number of homing runs from the threat area that directly targeted the obstacle edge (within 10 cm; correlation with post-removal escape targets: r=0.75, P=5×10^-5^; Fig. 4A-C). Two primary interpretations of these correlations are possible. The first is that routes are computed directly from a ‘cognitive map’: investigating the obstructed area updates the mouse’s internal map, which is reflected behaviorally in the mouse’s use of subgoals. If this were true, we would predict that: 1) investigating relevant features like the obstacle or its edge will also correlate with the subgoal memory; and 2) after obstacle removal, investigating the region where the obstacle used to be will suppress edge-vector escapes. Neither prediction matched the data. The amount of exploration near the obstacle or the obstacle edge was not correlated to subsequent escape target scores (correlation with distance moved around the obstacle: r= −0.09, P=0.7; with distance moved around the obstacle edge: r=0.06, P=0.8; Extended Data Fig. 6A). Furthermore, after obstacle removal, mice that densely sampled the empty center of the arena more did not execute different escape trajectories from mice that explored very little (correlation with distance moved around where the obstacle used to be: r= −0.12, P=0.6; with total post-removal exploration distance: r= −0.17, P=0.4; Extended Data Fig. 6A; see Extended Data Fig. 4A for examples of exploration during this period).

**Figure 4.**
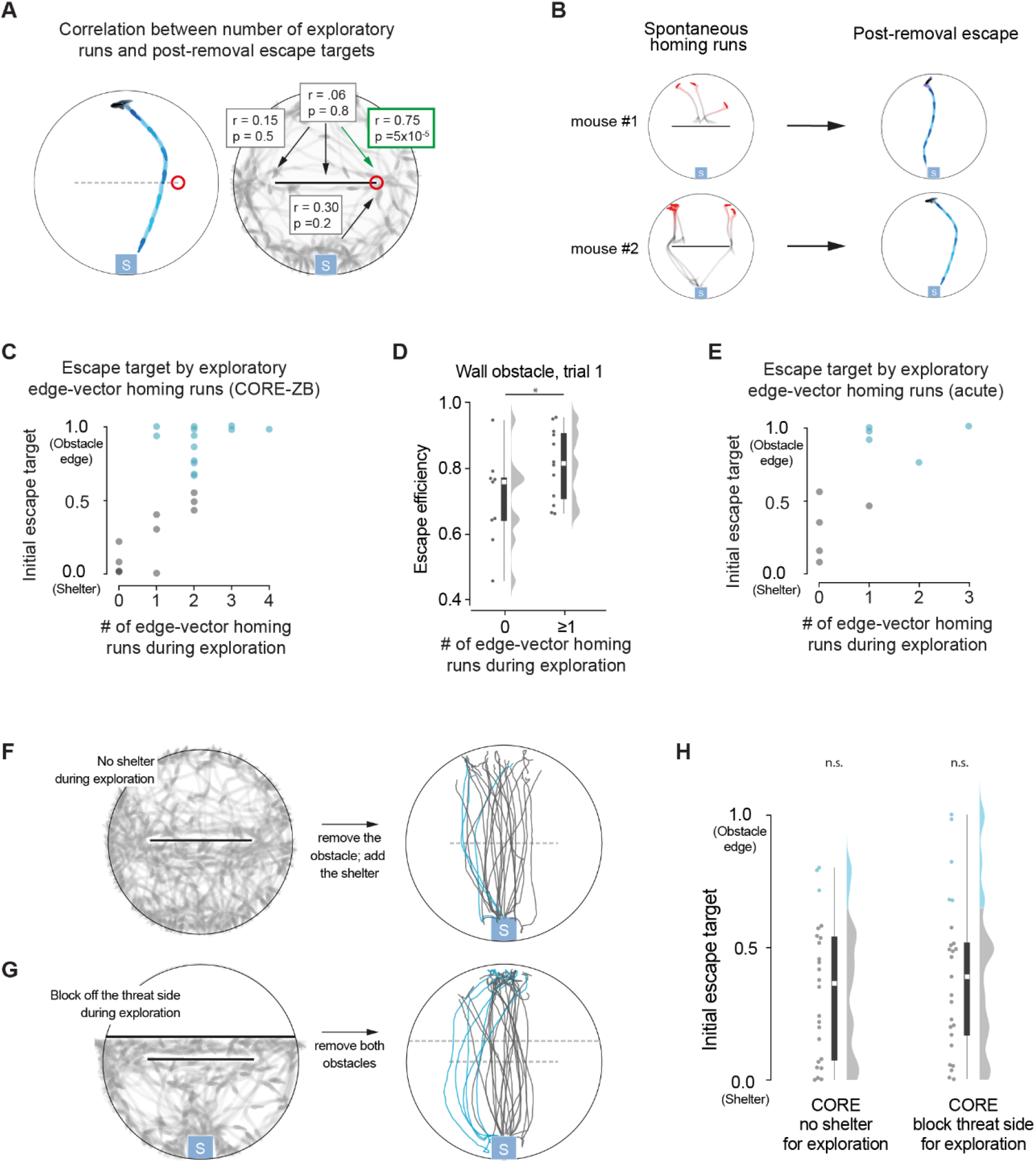
Mice memorize previously targeted subgoal locations. **(A)** Correlation of different running movements with the escape target score, in the CORE-ZB. These include homing runs from the threat area to different parts of the obstacle, as well as runs from the shelter area to the obstacle edge. Movements toward the same edge targeted in the escape (here, the right edge) are considered separately from movements toward the opposite edge (here, the left). Boxes show the correlation coefficients and respective p-values; significant correlations have green outlines. (B) Homing run history for two mice in the CORE-ZB, and subsequent escape trajectories. (C) Escape targets plotted against the number of spontaneous edge-vector homing runs during exploration in the CORE-ZB. As shown in panel A, these only include movements toward the same edge targeted during the escape. (D) Spatial efficiency of escapes on the first trial in the presence of an obstacle (same data as in Figure 1). As in previous panels, edge vectors only include movements to the edge targeted during the escape. **(E)** Escape targets plotted against the number of spontaneous edge-vector homing runs during exploration, for acute obstalcle removal on the first trial. (F) Chronic obstacle removal experiment without a shelter present during the 20-min exploration period. All subsequent escapes are shown on the right. N=24 escpaes, 10 mice. **(G)** Chronic obstacle removal experiment with an extra barrier blocking the threat area during the 20-min exploration period. All subsequent escapes are shown on the right. N=25 escapes, 10 mice. (H) Summary of escape targets in the two modified CORE-ZB experiments. Statistical test compares the proportion of edge-vector escapes to the open-field condition, as in Fig. 2B.

A second possibility is that learning occurs during the ‘practice’ edge-vector homings. In this case, we would predict that: 1) subgoals do not form in mice with zero edge-vector homings; and 2) the correlation with spontaneous homing runs would be specific to the edge targeted during escape (i.e., left vs. right) and to the direction taken during escapes (i.e., from the threat side to the shelter side). Both predictions were confirmed by the data. Every edge-vector escape following obstacle removal was preceded by at least one homing run targeting that same edge (100% of post-removal edge-vector escapes have ≥1 prior edge-vector run; greater than chance: P=0.02, *permutation test;* Fig. 4C). Second, escape targets in the CORE-ZB were not significantly correlated with homing runs from the threat area to the opposite edge (r=0.15, P=0.5), with homing-vector runs from the threat area to the middle of the obstacle (r=0.06, P=0.8), or with runs from the shelter area to the same obstacle edge (r=0.30, P=0.2; Fig. 4A).

#### Mice memorize previously encountered target locations as subgoals: experiments

Our analysis of the CORE-ZB suggests that executing edge-vector homings, rather than sampling the environment, could be the rate-limiting step in spontaneously learning subgoals. To further test this hypothesis, we first examined whether spontaneous homings explain escape routes in the obstacle-present condition. On the first trial with an obstacle, mice with prior edge-vector homings performed more efficient escapes than mice with none (median spatial efficiency with zero runs = 0.76; with one run = 0.82; P=0.04, *permutation test*; Fig. 4D; same data from Figure 1). As expected, this effect was specific to runs from the threat area to the side of the obstacle used during the escape (Extended Data Fig. 6B-D). Thus, subgoal memorization does appear to play an adaptive role when perception of the obstacle is still available. Next, we examined the acute obstacle removal experiment. We could not apply correlational analysis to the acute obstacle removal after three trials since this dataset had 100% edge-vector responses and 100% prior edge-vector homings. Thus, we performed a new experiment, removing the obstacle acutely on the first trial (10±1 minutes into the session, *mean±std*). Here, 50% of escapes took edge-vector paths (N = 10 escapes, 10 mice; Extended Data Fig. 6E). Among the variables examined – exploration in different parts of the platform and various running movements – only the number of runs from the threat area to the edge used in the escape was significantly correlated with escape targets (r=0.71, P=0.02; Fig. 4E; Extended Data Fig. 6F-G). Furthermore, 100% of edge-vector escapes were preceded by at least one edge-vector homing (greater than chance: P=0.02, *permutation test;* Fig. 4E).

Next, we tested the practice-homing hypothesis with two new experiments. First, we repeated the CORE-ZB but without a shelter during the exploration period (N=24 escapes, 10 mice; Fig. 4F). This gives the mouse opportunity to observe the platform and obstacle, but without performing homings. After 20 minutes, we added the shelter and removed the obstacle as soon as the mouse entered the shelter (median [IQR] time to enter shelter: 84 [39, 154] sec). Subsequent escapes did not exhibit the subgoal memory (13% edge-vector escapes; not more edge vectors than in the open field: P=0.4, *permutation test;* Fig. 4F, 4H). Second, we repeated the CORE-ZB with an extra barrier blocking off the threat side of the platform during the exploration period (N=25 escapes, 10 mice; Fig. 4G). This prevents long-range homings while allowing investigation of the obstacle. Only 1/10 mice targeted the edge location with scores close to 1.0, and post-removal escapes did not significantly differ from the open-field control (20% edge-vector escapes; not more edge vectors than in the open field: P=0.2, *permutation test;* Fig. 4G-H). Both experiments thus demonstrate a dissociation between investigating the obstacle and memorizing subgoals, and further support the hypothesis that subgoal locations are learned through practicing homings.

#### Spontaneous edge-vector runs are instinctive exploratory actions

It remains unclear what prompts spontaneous edge-vector homings in the first place. One possibility is that during practice homings, a cognitive map is used to compute efficient routes to shelter; once this happens, subgoals are tagged for later use during escapes. Another possibility is that mice are innately predisposed to run to salient obstacle edges. Our data support the latter option. Spontaneous edge-directed movement occurs most during the first few minutes of the session and occur equally with or without a shelter in the environment (Extended Data Fig. 7A-B). When the obstacle is a hole instead of a wall (Extended Data Fig. 1B), edge-directed movement occurs with the same, low frequency as in the open field (computed relative to the location where the obstacle edges would be if the obstacle were present; Extended Data Fig. 7B). Correspondingly, it takes twice as long for mice to perform predominantly edge-vector escapes in the presence of a hole obstacle (20% edge-vectors escapes on trial 2-3, 67% edge-vector escape on trial 6-7; Extended Data Fig. 7C-D).

### Subgoal learning also supports food-seeking routes

While subgoal memorization enhances spatial efficiency in a static environment, it can also generate unnecessarily roundabout routes past an obstacle that no longer exists. In fact, edge-vector escapes can persist over at least 20 minutes and 7 trials following obstacle removal (Extended Data Fig. 8A-B). We considered that subgoal memorization may be specific to escape behavior, as mice might sacrifice flexibility for the sake of quickly reacting to imminent threats. To test this, we performed an obstacle removal experiment in the context of a less urgent, reward-based task (open field control: N=32 reward runs, 6 mice; obstacle removal: N=34 reward runs, 6 mice). First, we trained food-deprived mice to approach and lick a reward port in response to a 10-kHz tone, which indicated the availability of condensed milk at the port. This took place across 5 sessions, in an operant conditioning box (Extended Data Fig. 9A-C). Next, we transported this task to the platforms previously used for escape behavior. The shelter was replaced by the reward port, and the threat stimulus was replaced by the 10-kHz tone. To start the session, mice were given 20 minutes in the open-field or obstructed environment. This included ~1 food-approach trial per minute with start points throughout the platform, to facilitate transferring the task to this new environment. After this point, mice successfully ran to the reward port during tone presentation on 85% of trials starting in the trigger zone (the same region as the threat zone), but with slower reaction times than escape (median [IQR] time to start running toward the goal for food-seeking = 1.5 [0.7, 3.5] sec; for threat response = 0.6 [0.4, 1.2] sec; P=0.005, *permutation test*). Next, we removed the obstacle and triggered food-approach trials (trials occurred 5±3 minutes after obstacle removal, *mean±std*). Similar to escape routes, a large proportion of paths in the obstacle-removal condition initially targeted the obstacle edge location (53% edge vectors; P=0.006 compared to 12% in the open field, *permutation test;* Fig. 5A-B; Supplementary Video 9).

**Figure 5.**
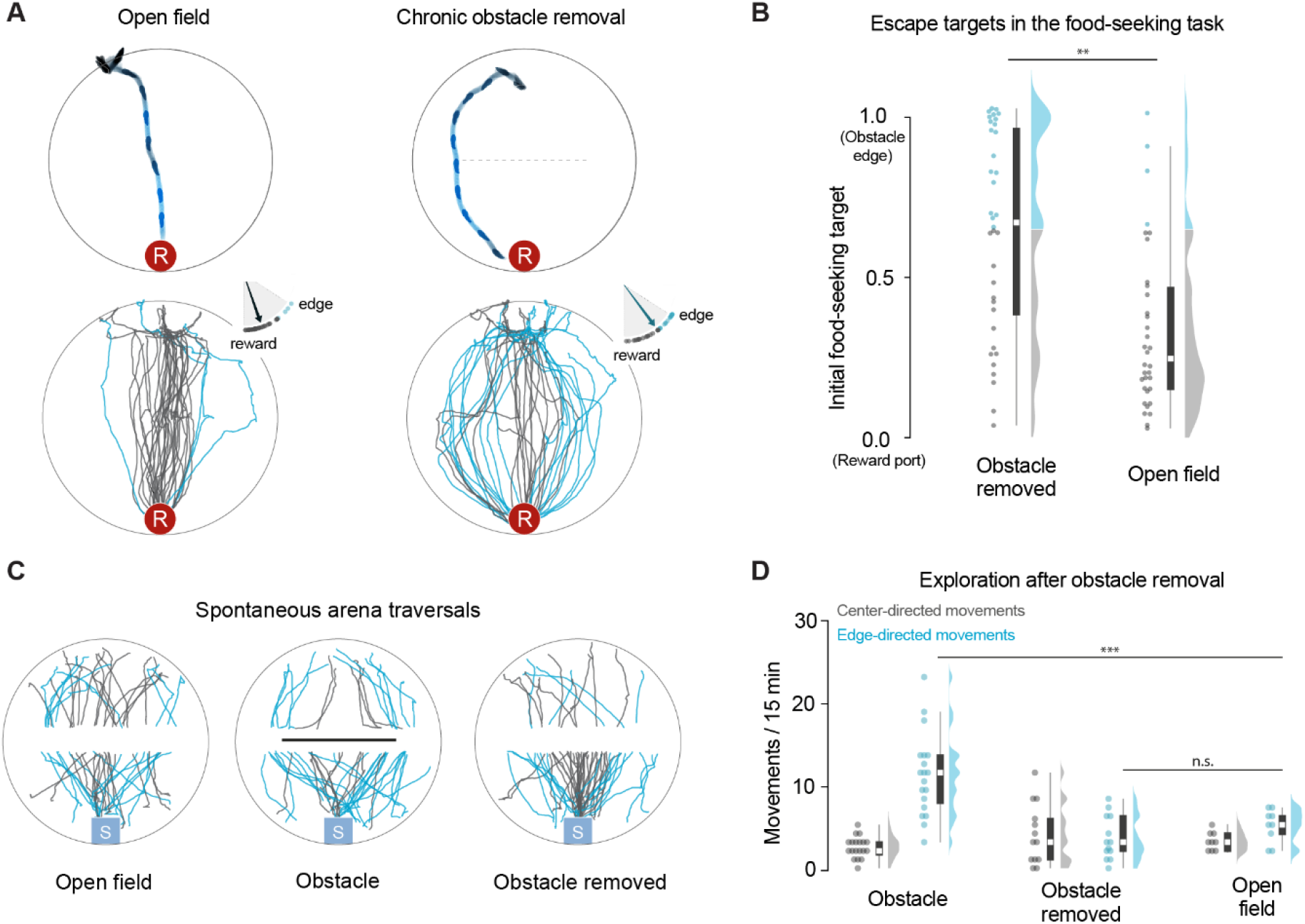
Experience with an obstacle changes food-seeking but not exploratory trajectories. **(A)** Food-approach paths in response to a 10-kHz tone associated with the availability of condensed milk at a reward port. An example trial is shown on top, and all paths are shown below. The red circle with ‘R’ represents the reward location, i.e. the metal spout with milk. The dotted line indicates the location of the obstacle during the initial 20-minute period. Open field: N=32 reward runs, 6 mice; chronic obstacle removal: N=34 reward runs, 6 mice. (B) Summary data for food-seeking paths, computed the same way as escape targets. The statistical test is a permutation test on the proportion of edge-vector paths. (C) Paths across the platform during spontaneous exploration in the escape experiments. All paths go from the ends of the platform toward the center. Conditions with more sessions are randomly downsampled so that the same number of paths is displayed for each condition. (D) The number of spontaneous center-directed and edge-directed movements during exploration. The statistical test is a permutation test on the number of edge-directed movements.

Finally, we tested whether experience with the obstacle induces a non-specific increase in edge-directed movement, as this could explain the apparent use of subgoal memorization across two distinct tasks. We compared spontaneous movements from the ends of the platform toward the center and obstacle edge locations. Exploration following obstacle removal were not enriched in edge-directed movements (number of edge-directed movements per 15 min: median after obstacle removal = 4; in the open field = 6; with the obstacle present =12; P=4×10^-5^, *permutation test on open field vs. obstacle;* P=0.7, *permutation test on open field vs. obstacle removed;* Fig. 5C-D). Subgoal memorization therefore reflects a strategy for goal-directed navigation rather than a general bias in how mice move around their environment following experience with an obstacle.

## Discussion

During their first few minutes in an obstructed environment, mice escaped to shelter by relying on their memory of the shelter location and their innate ability to negotiate barriers using vision and touch. These escape routes were spatially inefficient; they resembled obstacle avoidance in animals with lower cognitive capacities, such as toads, crabs, and ant colonies (Collett, 1982; Layne, 2003; McCreery et al., 2016). Over a single 20-minute session, however, mice began to exploit their aptitude for spatial memory. They increasingly targeted the obstacle edge directly and could do so even in complete darkness or after the obstacle had been removed. We found that this capacity relied on memorizing allocentric subgoal locations rather than egocentric turning movements, and our data further suggested that mice identified and memorized subgoals during spontaneous homing runs.

Previous work has shown that rodents use spatial memory to navigate to shelter in an open field (Alyan and Jander, 1994; Etienne et al., 1985; Vale et al., 2017). In such a simple environment, however, escape routes can be implemented by path integrating self-motion cues to keep track of a single vector to the shelter location – a one-step, egocentric process. With obstacles in the environment, a more advanced strategy is needed. Previous results in gerbils escaping in an obstructed environment suggested that spatial memory was employed to reach the shelter (Ellard and Eller, 2009), but their navigational strategy was unknown. Our results show that mice use subgoals in an allocentric reference frame. Several observations support this view. First, mice can accurately target the edge location minutes after the obstacle or the lights have been removed, which is not well explained by pure path integration. Second, escapes involved immediately orienting and running toward a subgoal ~50 cm away, which is not consistent with following odor trails or gradients (Liu et al., 2020; Wallace et al., 2002). Finally, repeating stereotyped turning movements or allocentric heading directions did not explain memory-guided escape paths in our assay; instead, mice consistently targeted the edge location. Future experiments on how escape routes transfer across multiple days, obstacles and tasks will help to specify the nature of this allocentric schema and how it is updated.

Traditional models of allocentric navigation involve three key elements: an internal map of the environment (located in the hippocampus and entorhinal cortex), a stored goal location, and a mental search for paths to the goal (Burgess et al., 1994; Edvardsen et al., 2020; Spiers and Gilbert, 2015; Stachenfeld et al., 2017). The limiting factor is the quality of the map. Finding efficient multi-step routes – be it through a tree-search algorithm (Edvardsen et al., 2020; Spiers and Gilbert, 2015), a map-partitioning algorithm (Stachenfeld et al., 2017), or warping around an ‘obstacle-to-avoid’ feature (Burgess et al., 1994) – can occur as soon as the map faithfully reflects the current environment. To build up this map, animals simply have to investigate unfamiliar or altered parts of the environment. The amount of exploratory movement thus matters for spatial learning, but movements’ intentions or directions do not (Burgess et al., 1994; cf. Schölkopf and Mallot, 1995). Our observations of escape routes in naïve mice do not support such views of allocentric learning. In our data, none of the following was sufficient to generate subgoals: 1) spending time exploring the obstacle; 2) running along the homing-vector path and then being blocked by the obstacle; 3) learning a subgoal at the other obstacle edge; 4) targeting the obstacle edge while running away from the shelter; 5) investigating the obstacle in the absence of a shelter; and 6) investigating the obstacle while the threat area was blocked off. Furthermore, investigating the formerly obstructed area following obstacle removal did not restore direct homing-vector responses.

The subgoal strategy does contain elements of classical map-based navigation: it is learned in all-or-none fashion and depends on a sense of allocentric space, i.e. a ‘map’; however, it also includes a component similar to taxon navigation, in which animals learn inflexible routes based on previous goal-directed movements (O’Keefe and Nadel, 1978). Hybrid strategies – combining rapid learning, inflexible routes, and special ‘learning movements’ – have been discovered before, as in the orientation flights of wasps (Collett, 1995). However, orientation flights entrain one-step routes to a visual beacon rather than multi-step routes to an obstructed goal. Our working model is that mice instinctively execute visually guided movements toward a salient wall edge; if this movement gives the mouse direct access to a subsequent goal (e.g., the shelter), then its target is memorized as a subgoal location. We hypothesize that a rapid, all-or-none learning rule works on practice homings, but further experiments should be done to test the causal role of this putative moment of insight.

Memorizing subgoals confers distinct survival advantages: it can drive escape routes with the optimality of map-based planning and the rapidity of instinctive responses. However, this strategy is less flexible than responding to sensation or updating maps. The steady persistence of ~50% biphasic escapes for tens of minutes after removing the obstacle was longer than expected, and it remains unclear how mice learn to reinstate the homing-vector response after obstacle removal. Responses to imminent predatory threats are known to favor quick reaction times at the expense of computational sophistication (Mobbs et al., 2020), and so this inflexible strategy could in principle be specific to defensive behavior. However, we found that it was also used in a less urgent food-seeking task. Thus, subgoals appear to be a building block for quickly learning spatial locations important for survival.

Subgoal learning bears some resemblance to hierarchical reinforcement learning, a technique in artificial intelligence for learning multi-step behaviors (Sutton et al., 1999). However, the learning process we have observed in mice does not fit cleanly into the dominant ‘model-free vs. model-based’ framework for reinforcement learning agents (cf. Spiers and Gilbert, 2015). Rather, it fuses action repetition with a model of space: mice discover a map of individual subgoals through targeted exploration and learning heuristics.

Finally, our results provide an alternative entry point to studying the neural mechanisms of spatial learning. Experiments with constrained behaviors and open-field environments have been crucial for the spatial-memory field; they uncovered the hippocampal formation’s key role in allocentric spatial memory (O’Keefe and Nadel, 1978), identified important activity dynamics in the hippocampus (Jadhav et al., 2012; O’Keefe and Recce, 1993; Wilson and McNaughton, 1994), and demonstrated the dorsal striatum’s involvement in repeating inflexible routes (Doeller et al., 2008; Packard et al., 1989). However, models of navigation based on these data remain limited in their applicability to real-world learning. To improve on this, future work will benefit from an understanding of spontaneous learning strategies and complex behaviors (Datta et al., 2019; Krakauer et al., 2017; Mobbs et al., 2018). Probing brain activity during spontaneous subgoal memorization presents one such opportunity for reconciling neural models with natural learning. This behavior’s rapid learning profile and reliable, stimulus-locked routes make it particularly tractable for testing theories of hippocampal and striatal functions and of the neural dynamics that underlie them.

## Experimental procedures

### Animals

All experiments were performed under the UK Animals (Scientific Procedures) Act of 1986 (PPL 70/7652) following local ethical approval. We used 158 singly housed (1 week), male, 8-12-week-old C57BL/6 mice during the light phase of the light cycle. For the main experiments, data come from the mice’s first-ever behavioral session. Table 1 describes all groups of mice used for this study, including the experimental conditions, any excluded mice, and re-use across experiments.

### Behavioral platforms

The main platform was an elevated white acrylic circular platform 92 cm in diameter (Extended Data Fig. 1A). The platform had a 51 cm x 1 cm hole in its center; through this hole, the obstacle (white acrylic, 50 cm long x 12.5 cm tall x 0.5 cm thick) could be raised (obstacle condition) or lowered (open field condition). For experiments in which the obstacle appears or disappears, this was done by digitally triggering a custom-made pneumatic tubing system (time to raise or lower the obstacle was ~100 ms; Supplementary Video 10). In the acute obstacle removal experiment, this was triggered simultaneously with the stimulus onset. In chronic obstacle removal experiments, this was triggered while the mouse was in the shelter. Obstacle removal makes a “whooshing sound” (63 dB measured at the shelter) and usually triggers a startle response. The presence or absence of this response was unrelated to subsequent expression or extinction of the subgoal memory. The hole obstacle consisted of a 50 cm long x 10 cm wide rectangular hole in the center of the platform (Extended Data Fig. 1B). The modified platform with two narrow corridors consisted of the original platform with the obstacle, plus six additional panels (schematic in Extended Data Fig. 5B). Four of these panels were 50 cm long x 12.5 cm tall x 0.5 cm thick, and two were 12.5 cm long x 12.5 cm tall x 0.5 cm thick. Together, they formed two corridors that were 50 cm long x 7.5 cm wide and were at 65° and 115 ° angles relative to the axis of the central obstacle. The interior panels forming the corridor were made of red acrylic so that the IR camera could see through them; all other panels were made of white acrylic. The platform in the sideways-moving-obstacle experiment was an elevated white acrylic square platform 80 cm x 80 cm. The obstacle (white acrylic, 70 cm long x 12.5 cm high x 0.5 cm thick) was manually pulled to the “short” condition (obstructing 48 cm) or to the “long” condition (obstructing 66 cm) while the mouse was in the shelter. The shelter was a 10 cm cube of transparent red acrylic (opaque to the mouse). It included a mouse-hole-shaped entrance at the front and additional 2.5 cm tall square of red acrylic on top in order to prevent the mice from climbing on top. The platform was surrounded by a black, square, plastic surrounding. A projector screen was located above the platform. The platform was illuminated with 4 infrared lights (S8100-45-A-IR, Fuloon). Experiments shown in Extended Data Fig. 3 and Supplementary Video 5 were performed in complete darkness (0.00 cd/m2 of visible light). At this light level, mice did not react to rapidly waving a hand in front of them, which is perceived as highly threatening when light is available. For all other experiments, light was projected onto the screen at 5.2 cd/m2 using a projector (PF1000U, LG). Note that the room was not totally sonically insulated and that neither the black surround nor the overhead illumination was circularly symmetric; these asymmetries could all provide spatial orientation cues. The platform and shelter were cleaned with 70% ethanol after each session.

### Escape behavior

Animals were given a 7-minute acclimation period during which they discovered the shelter. Stimuli were subsequently delivered when the mouse entered the threat zone (the back 20 cm, on the end opposite from the shelter) and was generally facing away from the shelter. Only stimuli delivered in this zone were analyzed. At least one minute was allowed in between trials. Threat stimuli were loud (87 dB), unexpected crashing sounds played from a speaker located 1m above the center of the platform. Sounds (“smashing” and “crackling fireplace”) were downloaded from soundbible.com. They were then edited using Audacity software such that they were 1.5 seconds long and continuously loud. Stimuli were alternated between the “smashing” sound and the “crackling” sound to prevent stimulus habituation. In some sessions (four with and four without an obstacle), we used an ultrasonic sweep stimulus (17-20 kHz, 3 sec). No difference in response between the stimuli was observed and therefore the data from these sessions were pooled. For each trial, the stimulus was triggered repeatedly until the mouse reached the shelter, for a maximum 9 seconds. Since escapes take longer with the hole obstacle and in the dark, stimuli in these conditions were played for up to 12 seconds. Stimulus responses were considered as escapes if the mouse reached the shelter within 12 seconds in the light or 18 seconds for the hole obstacle and for the dark. Stimulus delivery was controlled with software custom-written in LabVIEW (2015 64-bit, National Instruments). Stimuli were triggered manually, when the mouse had been in the threat zone (demarcated on the live video) for at least one second and was facing in approximately the opposite direction from the shelter. While manual stimulation could be a source of bias, we found that mice executing homing-vector vs. edge-vector escapes did not have significantly different starting positions or heading directions, limiting the impact that this bias could have (Extended Data Fig. 10A-C). We also found that the obstacle-removal experiments with differing results did not exhibit significantly different starting positions or orientations (Extended Data Fig. 10D). The sound was played from the PC, through an amplifier (TOPAZ AM10, Cambridge Audio) and speaker (L60, Pettersson). The audio signal was fed in parallel through a breakout board (BNC-2110, National Instruments) into a multifunction I/O board (PCIe-6353, National Instruments) and sampled at 10 KHz. To synchronize the audio and video, this signal was compared to the 30 Hz pulse triggering video frame acquisition, which was also fed as an input to the input/output board and sampled at 10 KHz. To verify correct synchronization, in most experiments the audio output cable was also fed in parallel to an infrared LED (850nm OLSON PowerStar IR LED), which flashed in synch with sound presentation. Mice varied in how many trials they performed in each experiment, due either to remaining in the shelter rather than entering the threat zone or not escaping in response to the stimulus (7% of trials). We thus limited analysis to the first three escapes in each condition (more than 50% of mice completed at least three escapes in all experiments).

### Food-seeking behavior

Mice were food restricted to 85% of their baseline weight. Training and pretraining were done in a 60cm x 15cm rectangular arena, with a shelter on one side and a reward port on the other side. The reward consisted of a 7-μL drop of condensed milk (diluted 1:1 with water) delivered through the spout. For pretraining, during which the mouse learned to associate the metal spout with reward, 100 drops of milk were manually triggered and then collected by the mouse, with a minimum interval of 1 minute between each drop. They were then trained in five, ~1-hour sessions to approach and lick a metal spout in response to a 9-second, 10-kHz, 72-dB tone. Tone stimuli were triggered manually once per minute. Licking the spout while the tone was on resulted in reward. After reward delivery, there was a 5-second refractory period; thus, mice could trigger at most two rewards during the 9-second tone. On the last two day of training, the tone duration was reduced to 4.5 seconds after 30 minutes. Licks were registered with a capacitive touch sensor (Adafruit MPR121), connected to a microcontroller board (Arduino Uno). The milk was delivered through a peristaltic pump (Campden Instruments 80204E), connected to the same microcontroller. For testing food-seeking paths, these mice had two sessions. The first session was in the platform with no obstacle, the shelter on one side, and a lick port on the opposite end of the platform. They received “practice trials” of tone and milk, initially mostly when they were already near the lick port. After 20 minutes, test trials were initiated when the mouse was on the opposite side from the lick port, and these data were used for analysis. The second session followed the same protocol. However, in this session the obstacle was initially present, and then was removed after 20 minutes while the mouse was in the shelter. Mice performed more trials than with the escape behavior, so here we examined trajectories from the first nine successful trials (greater than 50% of mice completed at least nine trials).

### Video tracking and visualization

Videos were acquired at 30 frames per second using an overhead camera (acA1300- 60gmNIR, Basler) with a near-infrared-selective filter. Video recording was performed with software custom-written in LabVIEW (2015 64-bit, National Instruments). Videos were then fisheye-distortion corrected, aligned onto a common coordinate framework, and visualized with custom Python code using the OpenCV library (github.com/BrancoLab/Common-Coordinate-Behaviour; Supplementary Video 1; Bradski, 2000). We used DeepLabCut (Mathis et al., 2018) to track the mouse from the video, after labelling 1500 frames with 13 body parts: snout, left eye, right eye, left ear, neck, right ear, left upper limb, upper back, right upper limb, left hind limb, lower back, right hind limb, tail base. Post-processing includes removing low-confidence tracking, using a median filter with a width of 7 frames, and applying an affine transformation to the tracked coordinates to match the common coordinate framework.

### Analysis

All analysis was done using custom software written in Python 3.6 (github.com/BrancoLab/escape-analysis) as well as open-source libraries, notably OpenCV and scikit-learn (Bradski, 2000; Pedregosa et al., 2011). The data that support the findings of this study are available from the corresponding authors upon request.

#### Calculating position, speed, and allocentric heading direction (Supplementary Video 11)

For analysis of trajectories and exploration, we used the center of the mouse. This was calculated as the average of all 13 points, which we found to be more stable and consistent than using any individual point. To determine the mouse’s speed for the color-coded visualizations, we smoothed the raw frame-by-frame speed with a gaussian filter (sigma = 3 frames = 0.1 sec). To calculate the mouse’s heading direction, we computed the vector between the center of the body (averaging the tail base, right hind limb, lower back, left hind limb, right upper limb, upper back, and left upper limb points) and the front of the body (averaging the left upper limb, upper back, and right upper limb points). We set the south direction (threat to shelter) to 0°, north (shelter to threat) as 180°, and east/west (left/right sides) as ±90°.

#### Quantification of escape targets

The initial escape target was computed by taking the vector from the mouse’s position at the escape initiation to its position when it is 10 cm in front of the obstacle. For the hole obstacle, this means 10 cm in front of the obstacle’s outer perimeter, rather than its center. We computed a target score where a vector aimed directly at the shelter received a value of 0; one aimed at either obstacle edge received a value of 1.0; a vector halfway between these would score 0.5; and a vector that points beyond the edge would receive a value greater than 1.0. The formula is: 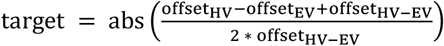, where offset_HV_ is the distance from the mouse to where the mouse would be if it took the homing vector, offsetEV is the distance from the mouse to where the mouse would be if it took the obstacle-edge vector, and offsetHV-EV is the distance from the homing-vector path to the obstacle-edge-vector path. Only the obstacle edge closest to the escape path was considered. Initial food-approach trajectories and spontaneous exploration trajectories were analyzed in the same manner. The threshold for classifying a trajectory as an edge vector was the 95^th^ percentile of escapes in the open-field condition (0.65), which is equivalent to the 10^th^ percentile of escapes on trial 3 with the obstacle. The rest were designated as homing vectors. Thus, we are slightly more conservative in labeling “edge vectors” (beyond the open field distribution with 95% confidence) than in labeling “homing vectors” (beyond the third obstacle escape distribution with 90% confidence). For examining the effect of experience on spontaneous exploration, we used a threshold of 0.5 to distinguish center-directed and edge-directed movements. In the narrow corridor experiment, a vector aimed at the left corridor opening received a value of 0, and a vector aimed at the left corridor opening received a value of 1.0.

#### Extraction of homing runs and escape initiation points

Homing runs are continuous turn-and-run movements from the threat side of the platform toward the shelter side. They are defined as a series of movements that continuously bring the mouse closer to the shelter or obstacle edges. They are extracted by: 1) computing the mouse’s “homing speed”, i.e. speed with respect to the shelter or obstacle edges with gaussian smoothing (σ=0.5 sec) and the mouse’s “shelter turning speed”, i.e. rate of change of heading direction with respect to the shelter; 2) identifying all frames in which the mouse has homing speed ≥ 10 cm/sec or is turning toward the shelter at an angular speed ≥ 90°/sec; 3) selecting all frames within 1 second of these frames to include individual frames that might be part of the same homing movement but do not meet the speed criteria; 4) rejecting all frames in which the mouse is not approaching or turning toward an edge or the shelter; 5) rejecting sequences that do not decrease the distance to the shelter by at least 30%. Each series of frames that meet these criteria represents one homing run. The homing runs we analyzed were those that started within 5 cm of the threat zone (the “threat area”) and ended with 5 cm of the central axis (i.e., along the obstacle). The homing target location is where a homing run crosses a line parallel to and 5 cm in front of the obstacle. Edge-vector targets occur when this location is within 10 cm to the left or right of the obstacle edge. **The escape initiation point** is defined as the beginning of a homing run that goes from inside the threat zone to outside of the threat zone following a threat stimulus. This is computed in the same way for spontaneous homings. In practice, the escape initiation point occurs when the mouse starts turning to run along the path that leads it out of the threat zone. We use this this criterion because it allows us to fairly compare spontaneous and stimulus-evoked homings, it correctly rejects initial post-stimulus movement bouts directed away from the shelter, and it correctly identifies the beginning of a turn-and-run movement as verified by manual inspection of the videos. Illustration of the escape initiation points for the main experiments are displayed in Extended Data Fig. 11. To characterize the urgency of the task based in the escape task vs. in the food task, we use the metric of the time until running toward the goal. This is when the mice cross a threshold on the homing speed of ≥20 cm/sec.

#### Quantification of turning angles

Turning angles that initiated homing runs and escapes were taken as the difference between the mouse’s heading direction at the start of the movement (the homing-run or escape initiation point) and the mouse’s heading direction after it had traveled 15 cm away from this start location. The start location is when the mouse starts turning toward and/or moving toward the shelter or obstacle edge (see previous subsection). Left turns were defined as negative, and right turns were defined as positive. For predicting escape targets from turn movements, we first extracted all homing runs from the mouse’s previous exploration experience. We then identified the 1-3 homing runs most similar to the escape, using three different similarity metrics: the most similar turn angle, the closest starting position, and closest initial heading direction. For each homing run-escape pair, we computed what the escape target would have been if the mouse had turned the same angle that it had turned during the homing run, i.e. if it had repeated the previous egocentric action. Finally, we performed a linear regression between the predicted targets (x) and the actual escape targets (y) to find the proportion of variance (R^2^) in escape targets predicted using this assumption that mice repeat previous egocentric turns. 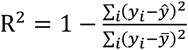, where 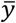 is the mean escape target and *ŷ* is the predicted escape target based on the linear regression. For the negative control, we disregarded the homing experience and instead predicted a random turn angle, and then extrapolated that angle to predict an escape target. We repeated this procedure 1000 times to get 1000 R^2^ values and took the mean R^2^.

#### Quantification of exploration

The time spent exploring was computed as the time spent at least 5 cm away from the shelter. The amount of exploration, or distance explored, was the time exploring multiplied by the mouse’s speed at each time point. Mice spent ~ 1/3 of the session in the shelter (IQR: 20-52% of the time). Spontaneous exploratory traversals are paths during exploration that start at either end of the platform (within 20 cm the end) and then reach within 10 cm of the central x-axis. Traversals that go along the boundary of the platform (i.e. within 10 cm of the outer perimeter) or take longer than 2 seconds (~10 cm/sec) were excluded from analysis, as these paths contained pausing and looping behavior, hindering the analysis of trajectories.

#### Statistical tests

For permutation tests, the test statistic is the group mean difference (e.g. in escape target or path efficiency). The condition of each mouse (e.g. open field vs obstacle) is randomly shuffled 10,000 times to generate a null distribution and a p-value. We used this test because it combines two distinct advantages: 1) Because the test statistic is the group mean, this test gives weight to each trial that a mouse performs rather than collapsing each animal’s data into one mean or improperly pooling trials (unlike the t-test or the Mann-Whitney test). 2) It is non-parametric and does not assume gaussian noise (unlike the repeated-measures ANOVA), in line with much of our data. Tests for differences in efficiency, reaction time, and initial escape conditions were two-tailed; tests for targets being biased toward previous experience were one-tailed. A different test statistic was used for the permutation test testing the significance of the result that 100% of memory-guided edge-vector escapes had at least one prior edge-vector homing. In this test, the p-value reflects the proportion of random subgroups of all trials that also score 100%. For the CORE-ZB (23 escapes, 13 edge-vector escapes), the p-value 0.02 indicates a 2% chance that every member of a random sub-group of 13/23 escapes has ≥1 prior edge-vector movement. The ANOVA was performed using the linear mixed effects model package in R, after removing outliers (z-score > 0.975). Fisher’s exact test was performed for cases with one trial per mouse and a categorical outcome. The Pearson correlation coefficient was used for correlation analyses. R2 values report the percent of the variance explained by the two variables’ linear relationship and is equivalent to the square of their Pearson correlation coefficient. The range illustrated in boxplots are limited from the first quartile minus 1.5 x IQR to the third quartile plus 1.5 x IQR.

## Acknowledgments

This work was funded by a Wellcome Senior Research Fellowship (214352/Z/18/Z) and by the Sainsbury Wellcome Centre Core Grant from the Gatsby Charitable Foundation and Wellcome (090843/F/09/Z) (T.B.), and the SWC PhD Programme (P.S. and S.O.). We thank members of the Branco lab and T. Mrsic-Flogel for discussions; J. Rapela for advice on statistical analysis; T. Mrsic-Flogel, T. Behrens, C. Barry, M. Stephenson-Jones, Y. Isogai, C. Clopath, Y.L. Tan, F. Claudi and R. Vale for comments on the manuscript; the SWC Neurobiological Research Facility and FabLabs for technical support; K. Betsios for programming the data acquisition software.

## Notes

### Competing Interest Statement

The authors have declared no competing interest.

### Summary of Updates

Updated statistical analyses, sample sizes, data presentation, and methods reporting. Reorganized the manuscript to clarify the flow of logic, particularly in the middle sections. Designed and executed four new experiments.

