## Extended Data Figures for "Mice learn multi-step routes by memorizing subgoal locations"

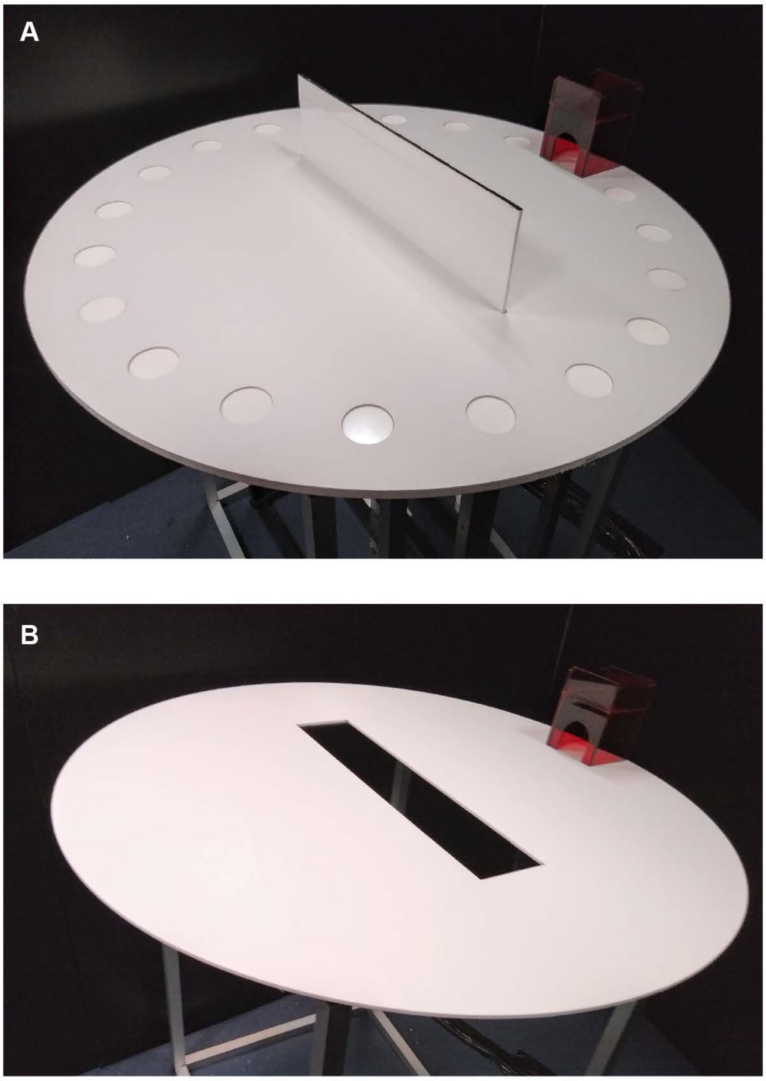


**Extended Data Figure 1 - Platforms with the wall obstacle and hole obstacle**

**(A)** Picture of the platform with the wall obstacle. The platform is 92 cm in diameter, and the wall obstacle is 50 cm long x 12.5 cm tall. The shelter is 10 cm long x 10 cm wide x 12.5 cm tall. It is made from red acrylic that is opaque to the mouse but transparent to red and infrared light. **(B)** Picture of the platform with the hole obstacle. The platform is 92 cm in diameter and has a 50 cm long x 10 cm wide x 1 m deep rectangular hole in the middle. Both plaforms are raised from the floor at a height of 1 m.

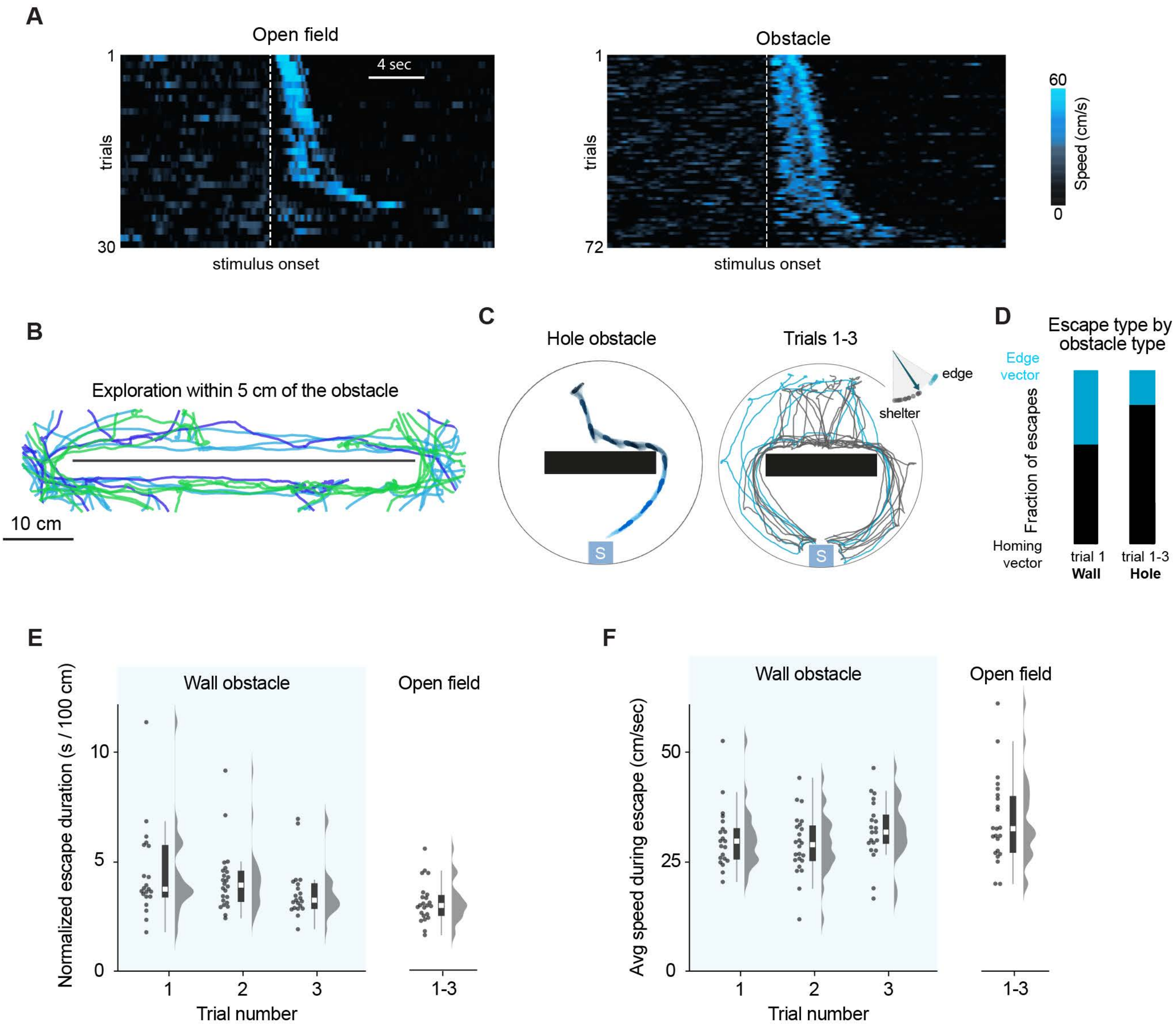


**Extended Data Figure 2 - Escapes in the presence of an obstacle**

**(A)** Speed profile for all escape trials (3 trials per mouse). Mice were stimulated with sound (dotted white line) while in the threat zone of the open field (N = 10 mice; left) or the platform with the obstacle (N = 24 mice; right). In the platform with the wall obstacle, the probability of threat-evoked escape to shelter was 93%, while the probability of escape in a no-stimulus control was 12%. Trials are sorted by shelter arrival time. **(B)** Exploration trajectories around the obstacle area (each color represents the movements of one mouse prior to trial 1). 3 randomly selected sessions are displayed. All mice approached and explored the area within 5 cm of the obstacle prior to the first escape trial (median time in this region: 37 sec, IQR: 29 - 45 seconds, minimum 14 seconds; median total distance explored in this region: 417 cm, IQR 259 - 493 cm, minimum 127 cm). **(C)** Example trial and escape trajectories for experiment with an unprotective hole obstacle insead of the wall obstacle (N = 23 escapes, 8 mice). Grey indicates homing-vector paths, and blue indicates obstacle-edge vector paths. The black rectangle represents the hole obstacle. **(D)** Summary of escape trajectories with the wall and hole obstacles. **(E)** Escape duration measured as the duration from escape initiation until reaching the shelter. To compare across conditions and trials, this value is normalized by the shortest possible path length to the shelter from the mouse's starting point. Each dot represents one escape. **(F)** Running speed during escape is the average speed of the mouse from escape initiation until reaching the shelter. There is a small effect of trial on running speed. $F(2, 37)=4.4$, $P=.02$, repeated measures ANOVA on trials 1-3.

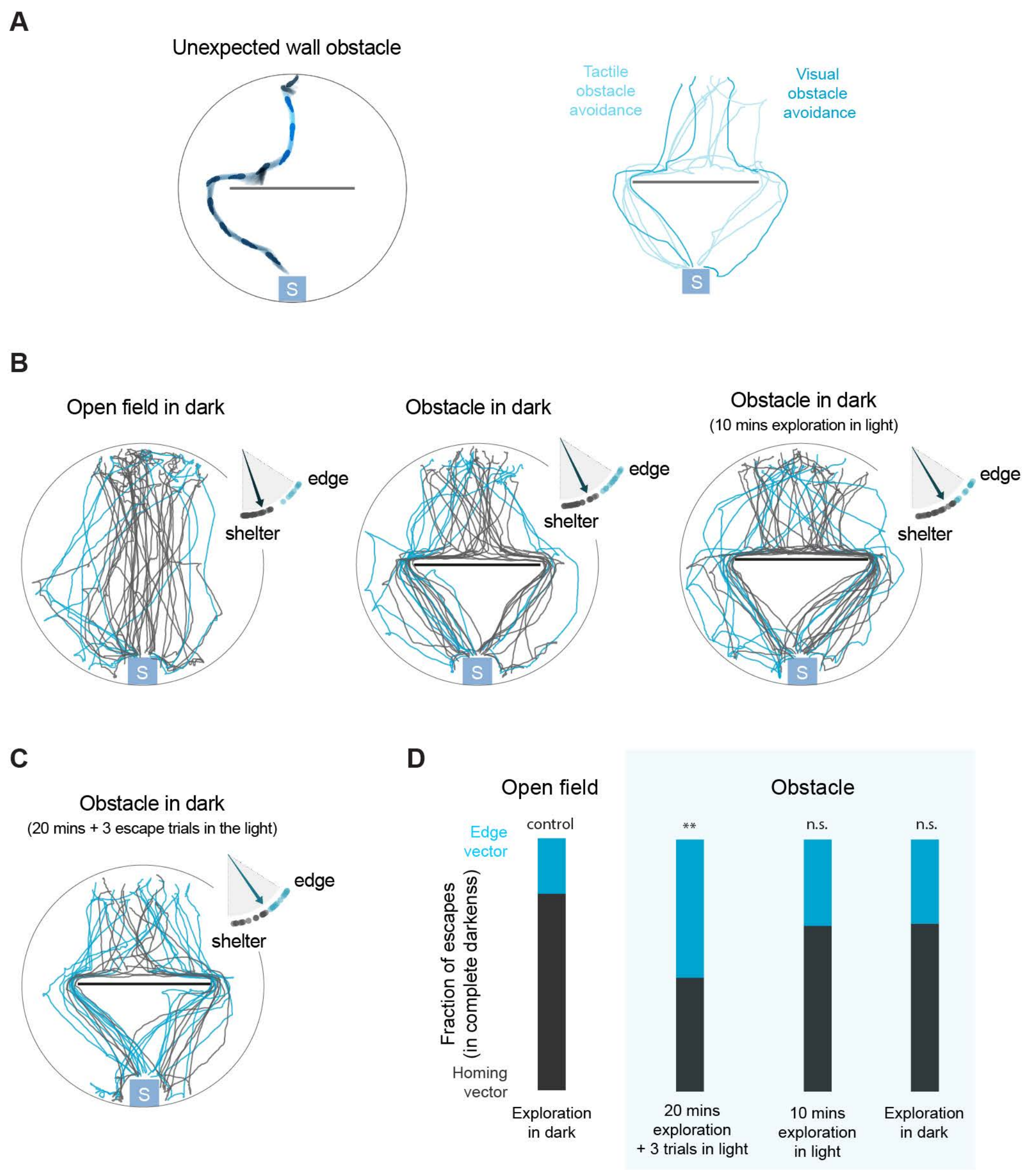


**Extended Data Figure 3 - The role of visual input in efficient obstacle avoidance**

**(A)** Example trial and all escape trajectories when the obstacle arises simultaneous with stimulus onset. This trial occurs after 20 minutes with three baseline escape trials in the open field. Putative visual avoidance occurs when the mouse turns toward an obstacle edge in the region between 5-10 cm away from the obstacle. Putative tactile obstacle avoidance occurs when the mouse turns toward an obstacle edge only after its head is already within 5 cm of the obstacle. N = 10 escapes, 10 mice **(B)** Escape trajectories for experiments in which naïve mice escape to the shelter in complete darkness (the four experiments in panels B and C are the only experiments in this paper performed in the dark). Dot-and-arrow plots display the distribution of escape targets in each condition. Open field: N = 41 escapes, 14 mice; obstacle: N = 33 escapes, 14 mice; obstacle + exploration in light: N = 33 escapes, 14 mice. **(C)** Mice with 20 minutes of experience in the light, including three escape trials. Even after removing all light, mice execute edge-vector responses. N = 32 escapes, 14 mice. **(D)** Summary of escape trajectories in complete darkness. ** P<0.01, permutation test on edge vectors in the dark, obstacle vs. open field.

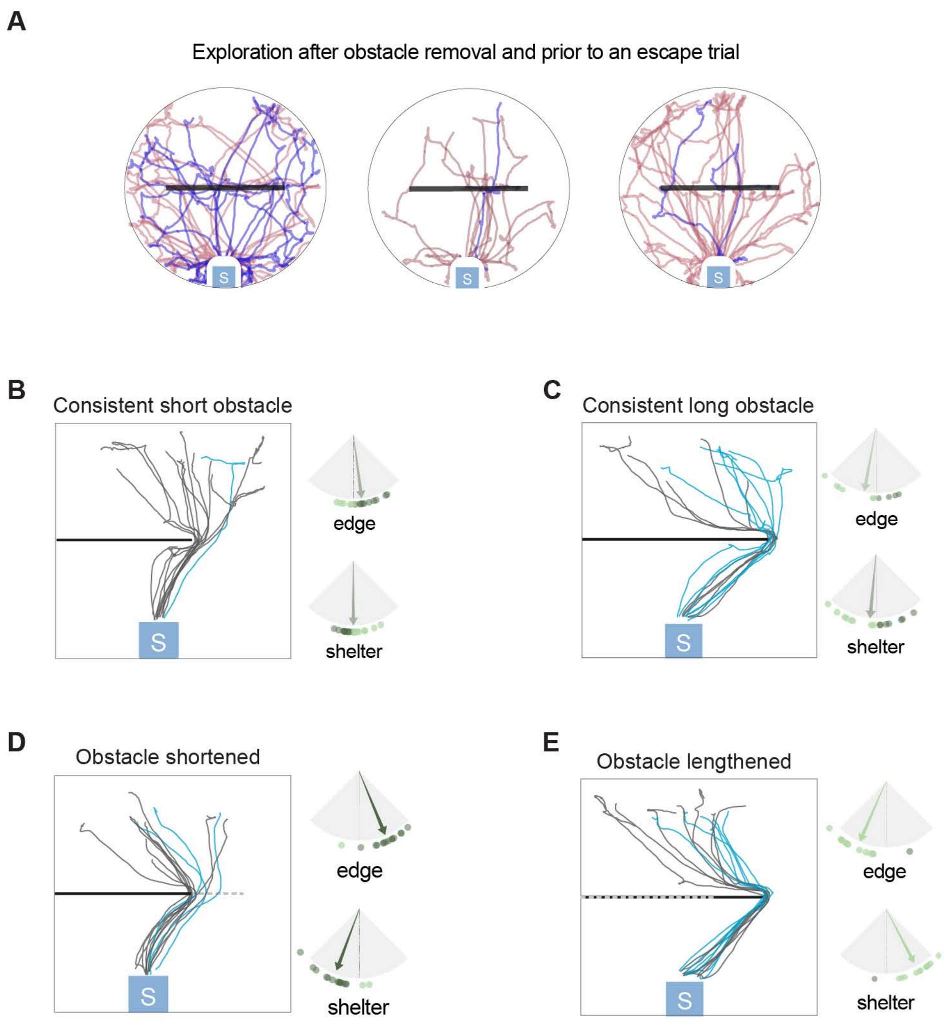


**Extended Data Figure 4 - Obstacle manipulation experiments**

**(A)** Chronic obstacle removal experiment. After the obstacle had been removed and prior to the escape trials, all mice previously visited the area where the obstacle used to be (black bar) during exploration. (This includes both the experiment with three baseline trials and the experiment with zero baseline trials.) Each colored trace represents the movements of one mouse after the obstacle was removed and prior to an escape trial. Six mice were randomly selected for visualization. **(B)** Escape trajectories in a control condition for the obstacle length-change experiment. In this condition, the obstacle is always short. N=15 escapes, 9 mice. Grey indicates trajectories targeting the shorter obstacle edge location, and blue indicates trajectories targeting the longer obstacle edge location. Arrow plots: dark green indicates overshooting the current edge or shelter position, and light green indicates undershooting (see Figure 2C). **(C)** Escape trajectories in a control condition for the obstacle length-change experimen. In this condition, the obstacle is always long. N=10 escapes, 8 mice. **(D)** Escape trajectories in an obstacle length-change experiment. The obstacle starts out long (dotted line) and is shortened after 20 minutes and 3 escape trials. N=14 escapes, 9 mice. **(E)** Escape trajectories in an obstacle length-change experiment. The obstacle starts out short (dotted line) and is lengthened after 20 minutes and 3 escape trials. N=13 escapes, 9 mice.

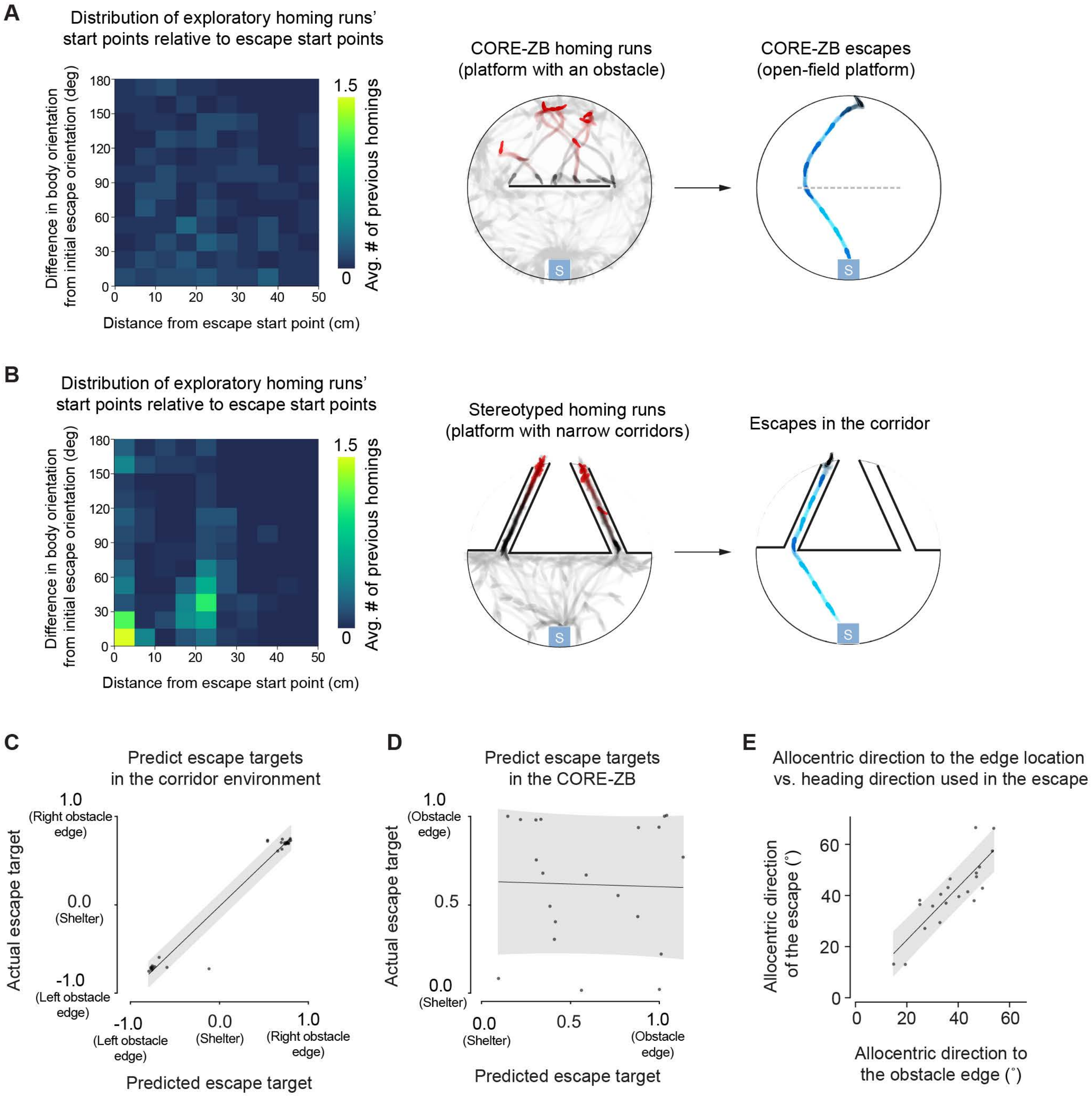


**Extended Data Figure 5 - Homing runs, turn angles, and heading directions**

**(A)** Left: histogram of homing runs' initial condition. This shows, for an average escape in the CORE-ZB, how many prior homing runs fell into different proximity bins. Each bin reflects proximity in both the position (x-axis) and body orientation (y-axis) of the homing's starting point. Right: example of homing runs extracted from exploration during the CORE-ZB's exploration period, and example of a subsequent escape in that experiment. **(B)** Experiment with narrow corridors that constrain movements during exploration and escape. Histogram and homing runs are computed in the same manner as in panel A. **(C)** Correlation between the predicted escape target (using the procedure illustrated in Fig. 3B) and the actual escape target. Here, escape targets are predicted using the homing run with the most similar turn angle to the escape turn angle. Data are from homings and escapes in the platform with narrow corridor. The correlation coefficient r=0.98; P=2x$10^{-22}$. This prediction thus yields an $R^2$ value of 0.97, which here corresponds to a mean absolute error of 0.025 in escape-target units. Note that we are using $R^2$ values as a measure of explained variance and not as a measure of goodness -of-fit of the linear relationship. Shaded area shows the prediction interval within 1 standard deviation. **(D)** Same analysis as in panel C, but with escapes and homings from the CORE-ZB. The correlation coefficient r= -0.03; P=0.9. This prediction thus yields an $R^2$ value of 0.0009, which corresponds to a mean absolute error of 0.38 in escape-target units. For comparison, predicting that every escape will be equal to the CORE-ZB's mean escape target generates a mean absolute error of 0.31. **(E)** Correlation between the allocentric heading direction required to target the obstacle edge location (x-axis) and the allocentric heading direction during escape (y-axis). The latter heading direction is measured when the mouse is 15 cm away from the escape initiation point (i.e., after the initial turn movement is complete). The vector from the center of the platform to the shelter (pointing south) is set as 0˚, and a vector pointing west or east is ±90˚. The absolute value of the heading direction is taken so that escapes toward the left and right edges can be considered together. Homing-vector escapes are not included. Data are from the two COREs of Fig. 2A-B.

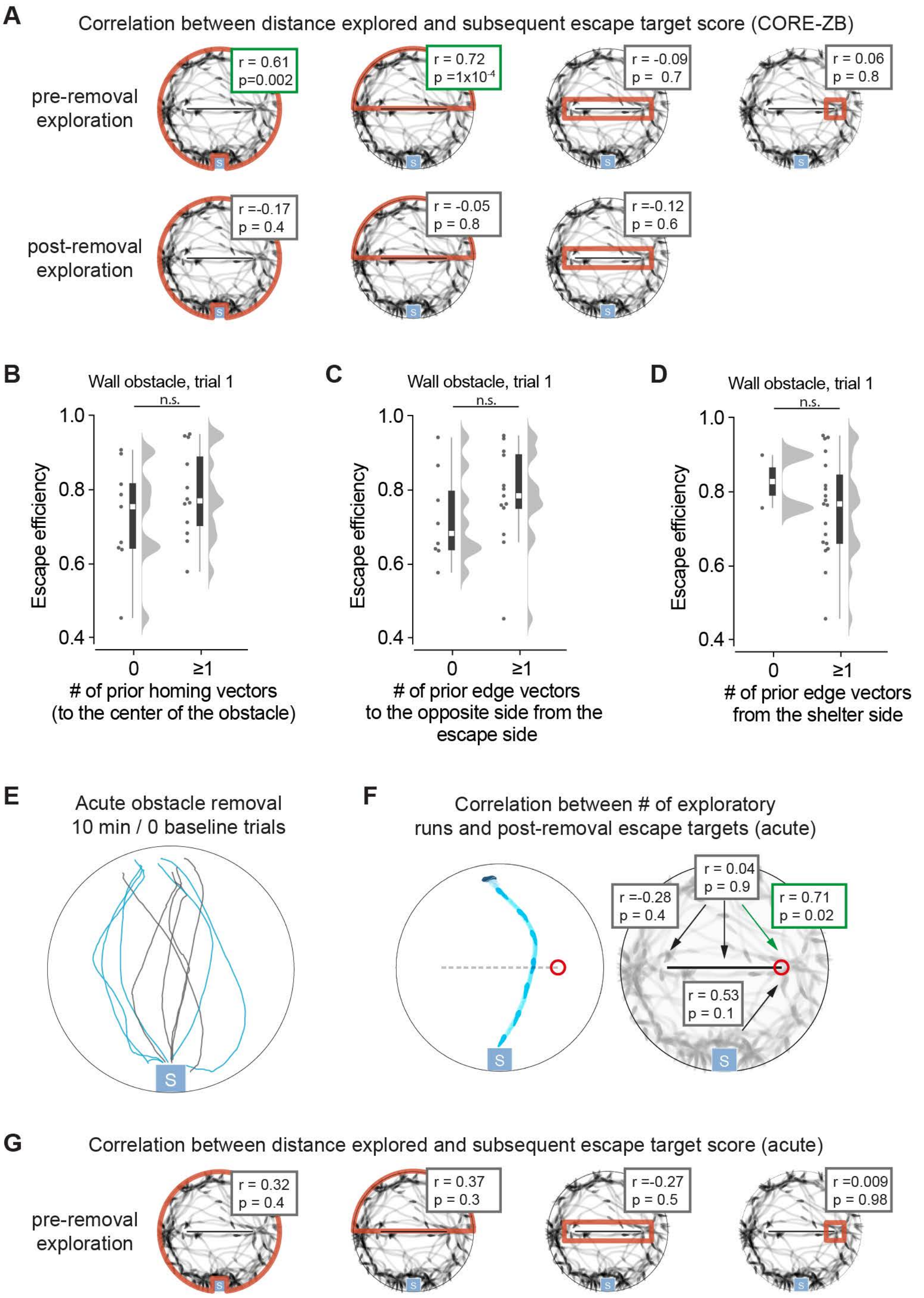


**Extended Data Figure 6 - Mice memorize previously targeted subgoal locations**

**(A)** Correlation between escape targets in the CORE-ZB and the amount of exploration in different sections of the platform. Red outlines indicate the section of the platform in which the distance explored is measured. For exploration near the obstacle edge, only the edge that was targeted during the escape (i.e., left vs. right) is considered. Boxes show the correlation coefficients and respective p-values; significant correlations have green outlines. **(B)** Spatial efficiency of escapes on the first trial in the presence of an obstacle (same data as in Figure 1). Here, runs from the threat area to the 10 cm in the center of the obstacle are considered. **(C)** Same as panel B, but here, runs from the threat area to the obstacle edge that was not used during the escape are considered. **(D)** Same as panel B and C, but for runs from the shetler area to the obstacle edge used during the escape. **(E)** Escapes from an experiment acutely removing the obstacle on the first trial, after 10 minutes of exploration. N=10 escapes, 10 mice. **(F)** Correlation of different running movements to escape target score, in the trial-1 acute removal experiment. These include homing runs from the threat area to different parts of the obstacle, as well as runs from the shelter area to the obstacle edge. Runs toward the same edge targeted in the escape (here, the right edge) are considered separately from runs toward the opposite edge (here, the left). **(G)** Correlation between escape targets in the trial-1 acute removal experiment and the amount of exploration in different sections of the platform. Post-removal exploration is not applicable in this experiment, since the obstacle is removed just before the escape begins.

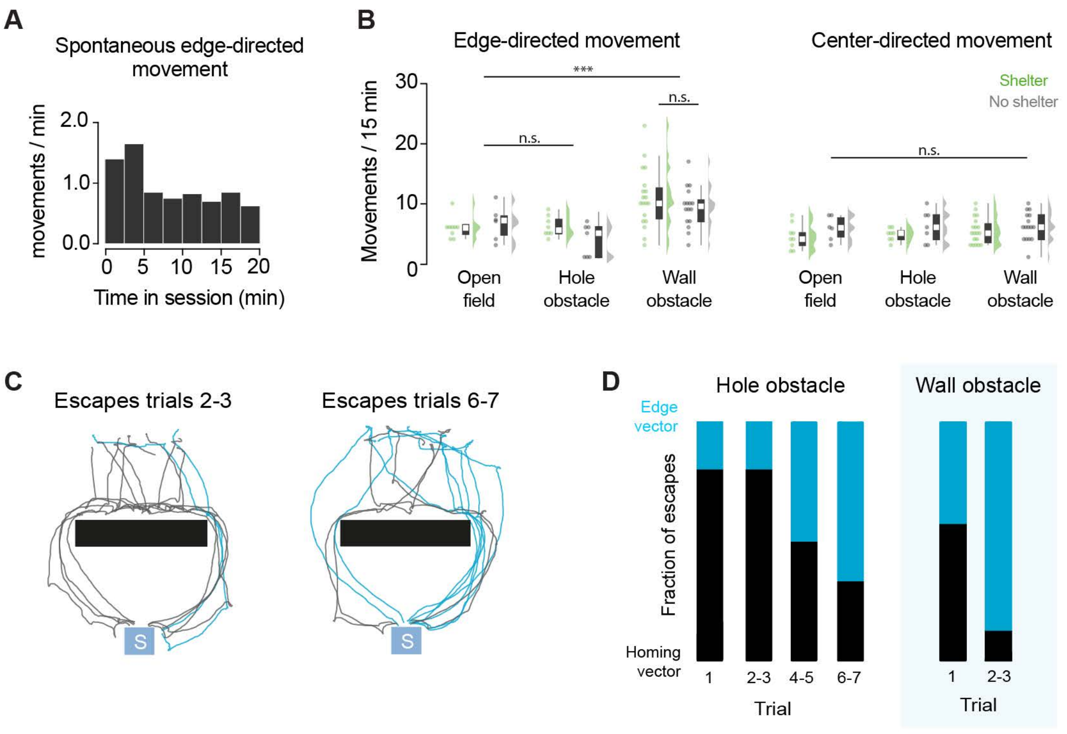


**Extended Data Figure 7 - Edge-directed movements in different environments**

**(A)** Frequency of spontaneous movements toward the obstacle edges (all sessions with an obstacle). **(B)** Frequency of edge-directed movements for different conditions. Number of edge-directed movements with the wall obstacle vs. open field: P=0.0005, permutation test. Open field (no shelter): N = 6 mice; hole obstacle (no shelter): N = 7 mice; wall obstacle (no shelter): N = 16 mice. For movements directed toward the center of the platform, there are no significant differences across conditions. Each dot is one session. **(C)** Escape trajectories for a hole obstacle (N=53 escapes, 8 mice). The black rectangle represents the hole obstacle. **(D)** Evolution of escape targets for increasing trial numbers with the hole osbtacle and the wall obstacle.

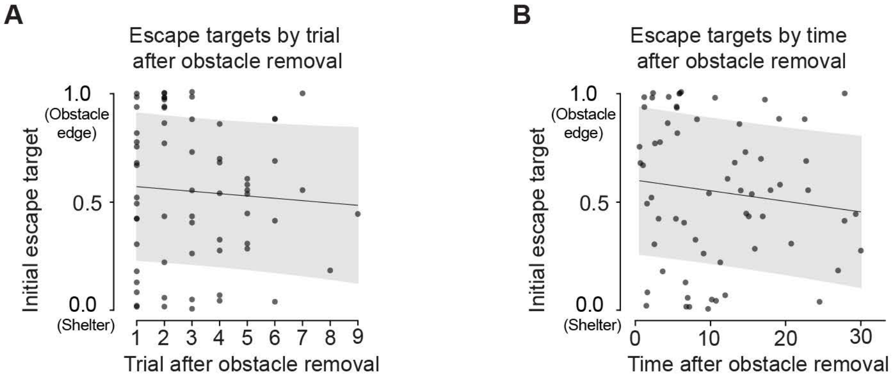


**Extended Data Figure 8 - Edge vectors persist over many trials and minutes after obstacle removal**

**(A)** Escape targets vs. trial number in the chronic obstacle removal experiments (3 baseline trials and zero baseline trials combined, and now including the minority of mice that performed >3 trials). For this plot, only successful escape trials are counted toward the trial number. The correlation between escape target and the number of stimulus-evoked escapes following obstacle removal is not significant: correlation coefficient r=-0.07, P=0.6. **(B)** Escape targets vs. time. The correlation between escape target and amount of time since obstacle removal is not significant: correlation coefficient r=-0.12, P=0.3. See Extended Data Fig. 6 for correlations between escape targets and the amount of post-removal exploration in various parts of the platform.

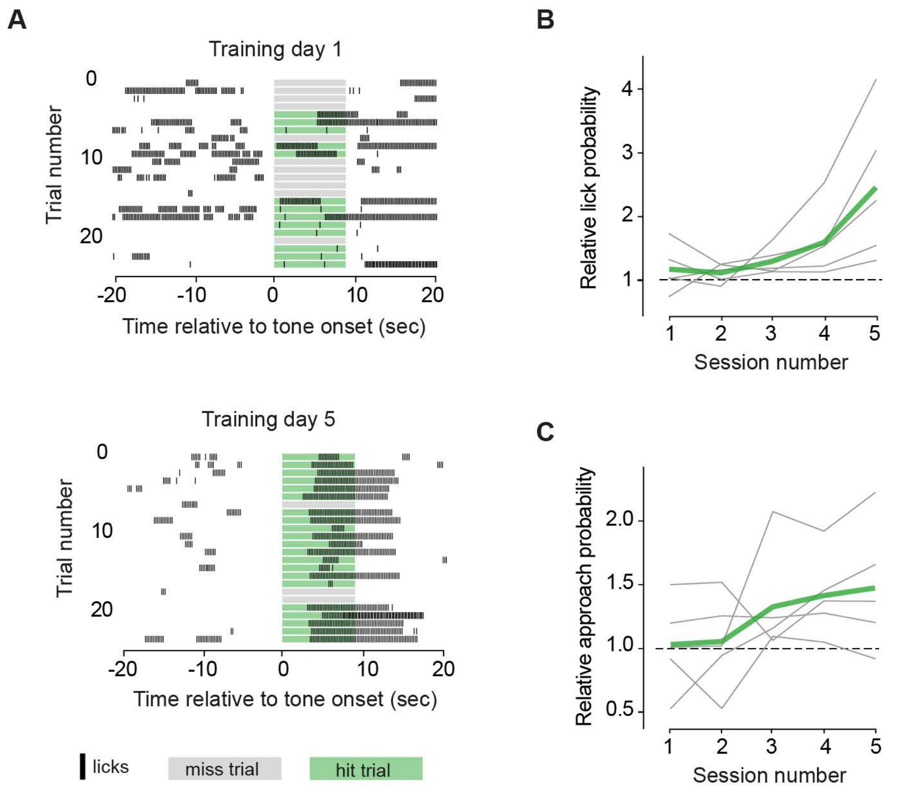


**Extended Data Figure 9 - Training mice to approach and lick a spout in response to a tone**

**(A)** Lick raster plots for an example mouse during the first (top) and the last training day (bottom). During food-approach training, a 9-second, 10-kHz tone is associated with the availability of condensed milk at a metal spout. For the lick raster plots, licks were plotted at 5 licks/sec when the sensor was tonically triggered by licking; this does not affect the quantifications in panels b-c. **(B)** Summary data for lick probability during training. Relative lick probability is the average probability of licking the spout within a 4.5-second window during the stimulus, divided by the lick probability during the 20 seconds before or after the stimulus. Mice lick the spout specifically in response to the tone on the fifth day of training (relative lick probability > 1, P=0.002, permutation test) but not on the first day (P=0.09, permutation test). **(C)** Summary data for reward-port approach probability during training. Relative approach probability is the average probability of moving from the back of the conditioning box to the side where the spout is located in response to the tone, divided by the approach probability at other random time points during the session. Mice approach reward specifically in response to the tone on the fifth day of training (relative approach probability > 1, P=0.02, permutation test) but not on the first day (P=0.41, permutation test). For **B** and **C**, gray lines are individual mice and green line is the mean. N=5 mice.

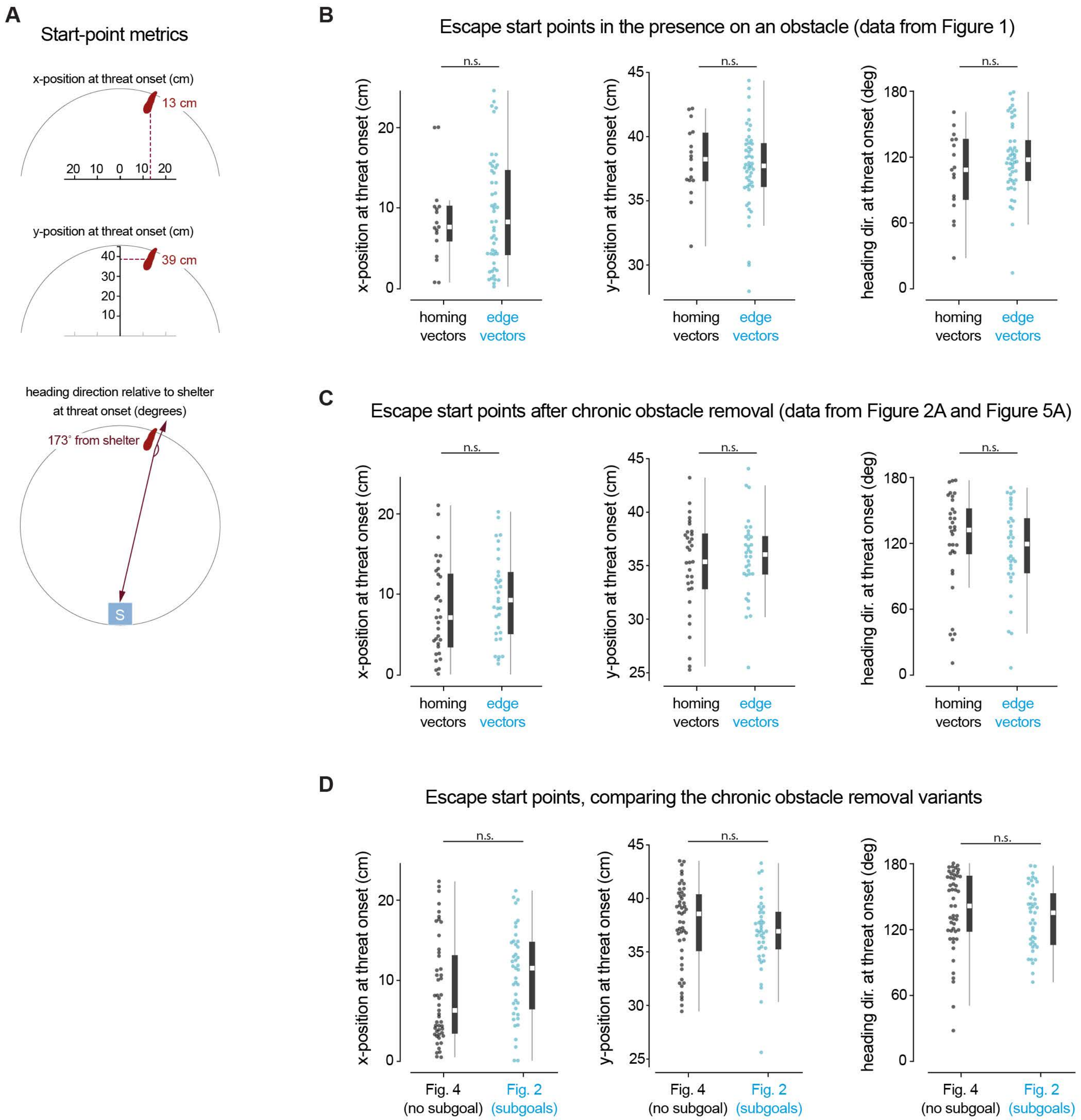


**Extended Data Figure 10 - Different responses and conditions are not associated with different start points**

**(A)** We examined three metrics to determine whether different initial positions upon threat onset were associated with different escape behavior. First, we examined the mouse's initial distance from the center of the platform along the horizontal axis (top). Second, we examined the mouse's distince from the center of the platform along the vertical axis (middle). Finally, we examined the mouse's body angle with respect to the shelter, i.e. the number of degrees it would need to turn in order to face the shelter. **(B)** Comparing start points for homing-vector responses vs. edge-vector responses in the obstacle condition. x-position: P=0.6; y-position: P=0.6; heading direction: P=0.2. **(C)** Comparing start points for homing-vector responses vs. edge-vector responses in the chronic obstacle removal experiments (CORE with zero baseline trials, CORE with 3 baseline trials, and CORE with the food seeking task). x-position: P=0.4; y-position: P=0.7; heading direction: P=0.6. **(D)** Comparing start points for the COREs that resulted in subgoal memory formation (Figure 2; CORE with zero baseline trials and CORE with three baseline trials) and the COREs that did not result in subgoal memory formation (Figure 4; CORE with no shelter during the exploration period and CORE with the threat side blocked off during exploration). x-position: P=0.06 ; y-position: P=0.3; heading direction: P=0.4.

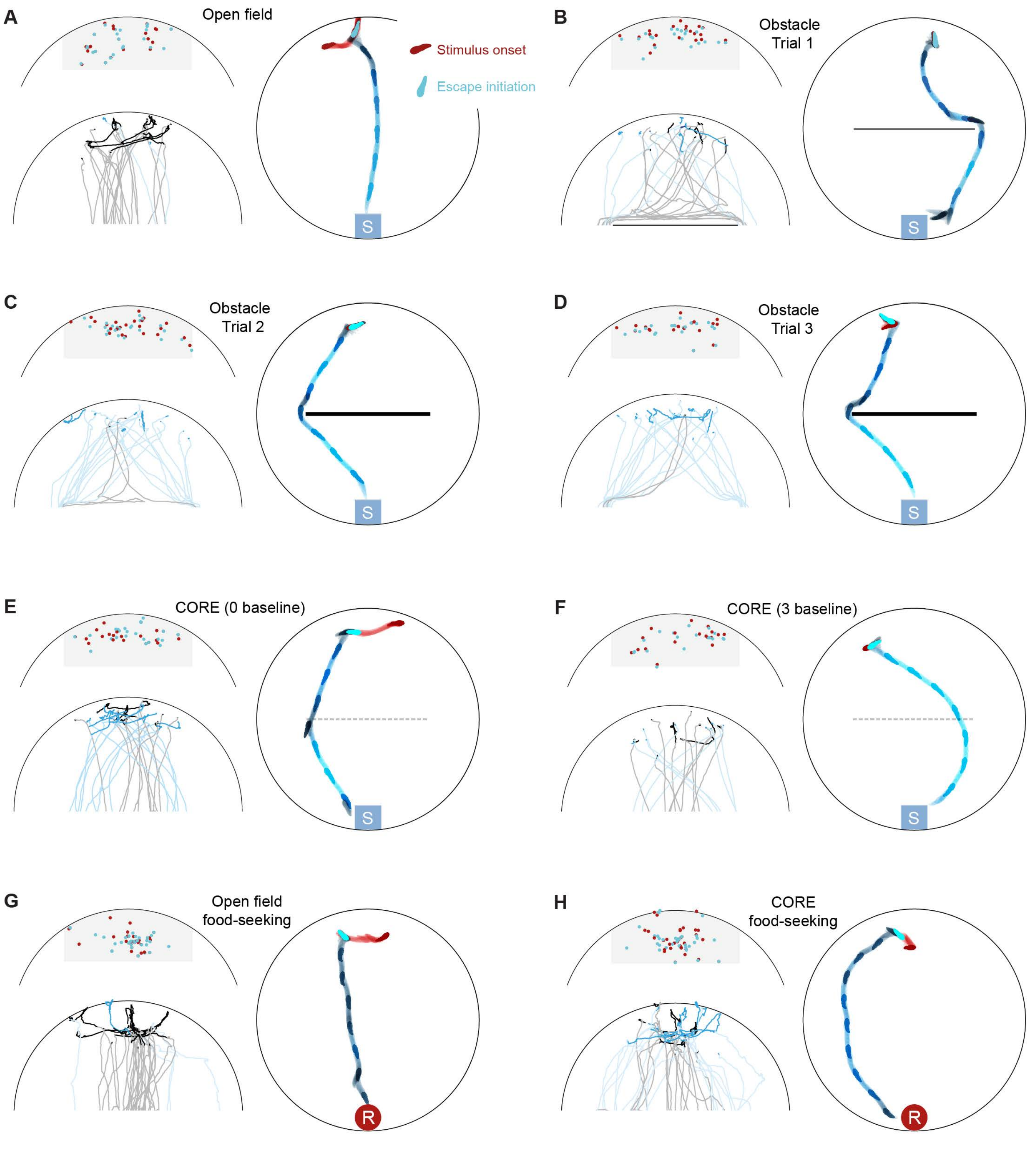


**Extended Data Figure 11 - Illustration of escape initiation points in the main experiments**

**(A)** Escapes in the open field. The escape initiation points mark the beginning of a turn-and-run movement from inside the threat zone (gray area) to outside the threat zone. This is computed the same way for escapes (shown here) and for spontaneous homing runs (Fig.3, Extended Data Fig. 5). Top left: Red dots show the mouse's position when the stimulus comes on for all trials, and blue dots show the mouse's position at the escape initiation point. Bottom left: Single escape trials are color-coded by trajectory type (homing-vector paths are black/gray, edge-vector paths are blue). Movements between the stimulus onset and the escape movement onset are shown in dark, bold traces. The rest of the escape is shown in a lighter hue. Right: example escape. The red mouse silhouettes mark the path between stimulus onset and escape initiation. The bright blue silhouette marks the escape initiation point, which is where the analysis of escape paths and turn angles begins. **(B)** Escapes with an obstacle (trial 1). **(C)** Escapes with an obstacle (trial 2). **(D)** Escapes with an obstacle (trial 3). **(E)** Escapes after chronic obstacle removal (zero baseline trials). **(F)** Escapes after chronic obstacle removal (three baseline trials). **(G)** Food-seeking trials in the open field. **(H)** Food-seeking trials after chronic obstacle removal.
