## Supplementary material for "Mice learn multi-step routes by memorizing subgoal locations": Table 1

| **Environment** | **Light** | **# of mice** | **Excluded - no threat zone entry** | **Previous stimulus experience** | **Time before trial 1** | **Data location** |
| --- | --- | --- | --- | --- | --- | --- |
| Open field | On | 10 | 0 | None | 10 min | F1, EDF 2, F5, EDF 10 |
| Obstacle | On | 24 | 0 | None | 10 min | F1, EDF 2, F4, EDF 6, F5, EDF 10 |
| Hole obstacle | On | 8 | 0 | None | 10 min | EDF 2, EDF 7 |
| Obstacle rises acutely | On | 10 | 5 | 3 trials (Open field) | 20 min | EDF 3 |
| Obstacle rises acutely | On | 9 | 4 | 1 session + 3 trials (Open field) | 20 min | EDF 3 |
| Open field | Dark | 14 | 0 | None | 10 min | EDF 3 |
| Obstacle | Dark | 14 | 0 | None | 10 min | EDF 3 |
| Obstacle | 10 min on then dark | 14 | 0 | None | 10 min | EDF 3 |
| Obstacle | 20 min on then dark | 14 | 0 | 3 trials (Obstacle) | 20 min | EDF 3 |
| Acute obstacle removal trial 4 | On | 10 | 2 | 3 trials (Obstacle) | 20 min | F2 |
| Chronic obstacle removal (CORE) #1 (CORE-3B) | On | 10 | 2 | 3 trials (Obstacle) +  1 trial (Acute removal) | 20 min | F2, F4A, EDF 8, F5, EDF 10 |
| CORE #2 (zero baseline trials; CORE-ZB) | On | 10 | 0 | None | 20 min | F2, F3, EDF 5, F4, EDF 6, EDF 8, F5, EDF 10 |
| Square w/ short obstacle | On | 12 | 3 | 1 session | 10 min | F2, EDF 4 |
| Square w/ long obstacle | On | 12 | 4 | 1 session | 10 min | F2, EDF 4 |
| Square w/ lengthened obstacle | On | 9 | 0 | 1 session + 3 trials (long obstacle) | 20 min | F2, EDF 4 |
| Square w/ shortened obstacle | On | 12 | 3 | 1 session + 3 trials (short obstacle) | 20 min | F2, EDF 4 |
| Narrow corridors | On | 10 | 0 | 1 session | 20 min | F3, EDF 5 |
| CORE #3 (additional short barrier) | On | 10 | 0 | None | 20 min | F3 |
| CORE #4 (move the shelter) | On | 10 | 0 | None | 20 min | F3 |
| Acute obstacle removal trial 1 | On | 10 | 0 | None | 10 min | F4, EDF 6 |
| CORE #5 (no shelter at first) | On | 10 | 0 | None | 20 min | F4, EDF 7 |
| CORE #6 (additional long barrier) | On | 10 | 0 | None | 20 min | F4 |
| Open field (no shelter) | On | 6 | 0 | 1 session | n/a | EDF 7 |
| Obstacle (no shelter) | On | 6 | 0 | 1 session | n/a | EDF 7 |
| Hole obstacle (no shelter) | On | 7 | 0 | 1 session | n/a | EDF 7 |
| Open field (food seeking) | On | 6 | 0 | 5 training sessions (conditioning box) | 20 min | F5, EDF 9, EDF 10 |
| CORE #7 (food seeking) | On | 6 | 0 | training + 1 session (open field food) | 20 min | F5, EDF 9, EDF 10 |

**Table 1. All groups of mice and all experimental conditions.** Mice that were excluded generally stayed inside of the shelter instead of exploring. This is more common in mice with more previous experience. Previous stimulus experience refers to previous trials and sessions from an experiment higher up on the list. For example, “3 trials (Obstacle)” come from the “Obstacle” condition listed higher up in the table.
